# ATPase activity of DFCP1 controls selective autophagy

**DOI:** 10.1101/2022.02.24.481614

**Authors:** Viola Nähse, Camilla Raiborg, Kia Wee Tan, Sissel Mørk, Maria Lyngaas Torgersen, Eva Maria Wenzel, Mireia Nager, Veijo T. Salo, Terje Johansen, Elina Ikonen, Kay Oliver Schink, Harald Stenmark

## Abstract

Cellular homeostasis is governed by removal of damaged organelles and protein aggregates by selective autophagy mediated by cargo adaptors such as p62/SQSTM1. Autophagosomes can assemble in specialized cup-shaped regions of the endoplasmic reticulum (ER) known as omegasomes, which are characterized by the presence of the ER protein DFCP1/ZFYVE1. The function of DFCP1 is unknown, as are the mechanisms of omegasome formation and constriction. Here, we demonstrate that DFCP1 is an ATPase that dimerizes in an ATP-dependent fashion. Whereas depletion of DFCP1 had a minor effect on bulk autophagic flux, DFCP1 was required to maintain the autophagic flux of p62 under both fed and starved conditions, and this was dependent on its ability to bind and hydrolyse ATP. While DFCP1 mutants defective in ATP binding or hydrolysis localized to forming omegasomes, these omegasomes failed to constrict. Consequently, the release of nascent autophagosomes from omegasomes was markedly delayed. DFCP1 was found to associate with ubiquitinated proteins, and degradation of ubiquitinated cargoes such as protein aggregates, mitochondria and micronuclei was strongly inhibited when DFCP1 was knocked out or mutated. Thus, DFCP1 mediates ATPase-driven constriction of omegasomes to release autophagosomes for selective autophagy.

## Introduction

Autophagy is a cellular degradation mechanism that is critical to maintain cellular homeostasis – either by recycling of bulk cytoplasmic components to produce energy in times of starvation or by degradation of potentially harmful cytoplasmic objects. During autophagy, cargo is engulfed by a double membrane, the phagophore. After sealing of the phagophore, the autophagosome – containing the engulfed cargo – fuses with a lysosome for degradation of its content ^1^.

The ER is considered the major site of autophagosome formation ^2 3^. A major regulator of autophagy is the class III phosphatidylinositol 3-kinase complex (PI3K-III), which is activated by pro-autophagic cues to generate phosphatidylinositol 3-phosphate (PtdIns3P) at specialized regions of the ER. These PtdIns3P-rich ER-subdomains serve as platforms for autophagosome assembly. They form characteristic cup-like structures, which are called ‘omegasomes’ ^4 5^. Within the omegasome cup, a double membrane, the phagophore, is formed, which extends and engulfs the cargo. During this process, it forms membrane contact sites with the ER, which likely provides the membrane for the extension of the phagophore ^5 6^. Once the phagophore has engulfed the cargo, it closes into an autophagosome and dissociates from the ER ^5 7^.

In line with the requirement for PtdIns3P, several PtdIns3P binding proteins are important for autophagosome formation and closure. During early phases of the phagophore formation, two groups of PtdIns3P binding proteins are recruited. Proteins of the WIPI (WD-repeat protein interacting with phosphoinositides) family ^8^ are necessary for lipid conjugation and activation of the phagophore-forming protein LC3B ^9, 10^. The other PtdIns3P-binding protein, DFCP1 (Double FYVE containing protein) ^4^ is recruited to omegasomes. While DFCP1 has been widely used as an early autophagy reporter, its cellular functions are largely unknown, and its role during autophagosome formation is not understood.

Here, we show that DFCP1 is an ATPase that dimerises in an ATP-dependent fashion and that it accumulates with ubiquitinated proteins at specific ER sites. We further demonstrate that the ability of DFCP1 to bind and hydrolyse ATP is required for omegasome constriction and selective autophagy.

## Results

### DFCP1 binds and hydrolyses ATP

The main structural features of DFCP1 are two C-terminal FYVE domains, which, in conjunction with an ER-targeting region, bind to PtdIns3P on ER domains and by this marks the sites of autophagosome formation ^4^ (Fig.1a). In contrast, the N-terminus of DFCP1 does not carry any characterized domains, but we found it interesting that it contains a P-Loop motif ^4^. A search in the Interpro databases revealed that the DFCP1 domain structure, including the P-loop, the ER-binding domain and the two FYVE domains, is an ancient architecture which evolved more than 500 million years ago, with homologues in most metazoan phyla (Extended Data Fig.1a,b). P-loops interact with phosphorylated nucleotides, but no function of the N-terminal domain has so far been described for DFCP1.

**Fig.1:**
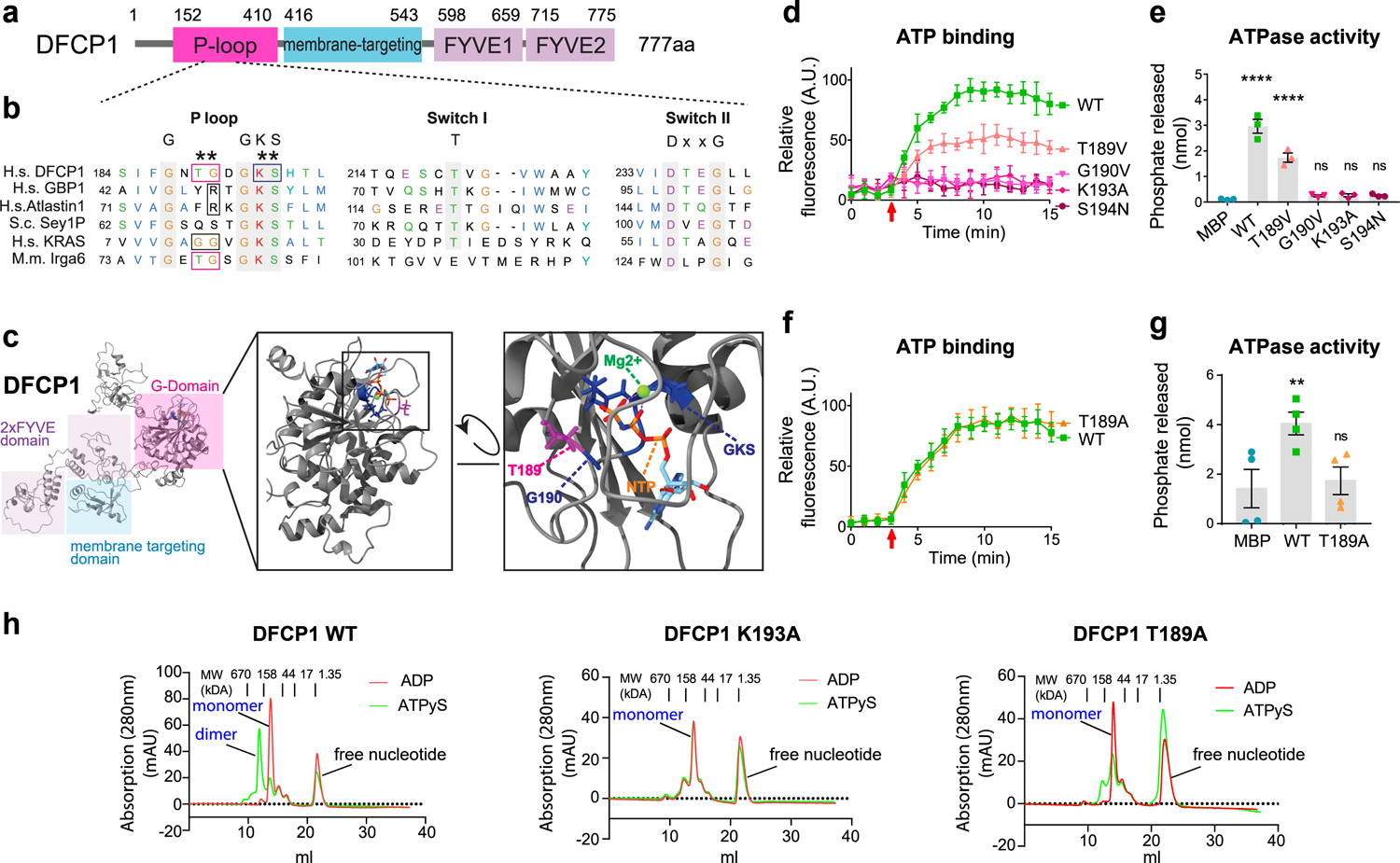
DFCP1 has structural similarities to ATPases and GTPases and binds and hydrolyses ATP. (a) Domain structure of DFCP1. DFCP1 contains an unstructured N-terminal domain, a putative P-loop nucleotide binding domain, a membrane binding domain and two FYVE domains. (b) Sequence alignments of the P-loop, switch I and switch II regions of DFCP1, GBP1, Atlastin1, Sey1P and KRAS. Boxes highlight residues critical for nucleotide hydrolysis in GBP1, Atlastin1 and KRAS. Asterisk indicates DFCP1 residues picked for mutational analysis. (c) AlphaFold-generated model of full-length DFCP1. Shaded regions indicate functional domains. Candidates for amino acids required for nucleotide binding (GKS, residues 192-194) are indicated in blue, and amino acids at the same position as the catalytic site of GBP1 (T189, G190) and KRAS (G12, G13) are indicated in magenta. The nucleotide (GDP) and Mg2+ ion were extracted from GBP1 (PDB ID: 1f5n) after structural alignment. (d) DFCP1 binds to ATP and identification of binding mutants. DFCP1 WT and mutants were incubated with mantATP (arrow), and nucleotide binding was measured by the incorporation of fluorescent mantATP. Curves are normalized to MBP background fluorescence. Bars: mean ± SD, 3 experiments. (e) Measurement of phosphate release by nucleotide-binding DFCP1 mutations. Purified DFCP1 and DFCP1 point mutations (10 µM) were incubated in the presence of ATP and the release of free phosphate was measured. Bars: mean ± SEM, 3 experiments. Statistics: One-way Anova followed by Dunnett’s multiple comparisons test, comparing against MBP. (f) Characterization of ATP-hydrolysis impaired DFCP1 point mutation. Purified DFCP1 and DFCP1 T189A were incubated in the presence of mantATP (arrow) and the binding of fluorescent mantATP was measured. Bars: mean ± SEM, 4 experiments. (g) DFCP1 T189A is unable to hydrolyze ATP. DFCP1 WT and DFCP1 T189A were incubated with ATP and the release of free phosphate was measured. Bars: mean ±SEM, 4 experiments. Statistics: Ordinary One-way Anova followed by Dunnett’s multiple comparisons test, comparing against MBP. (h) Purified His6-MBP-DFCP1 WT or mutant (aa 1-410) were loaded with ADP or ATPγS and the apparent molecular weight was determined by size exclusion chromatography. Shown are the elution profiles in response to ADP or ATPγS loading. Relative molecular weights were determined by inclusion of a standard (BioRad Gel filtration standard). The peak eluting at the higher molecular weight corresponds to a dimer, whereas the lower molecular weight corresponds to a monomer. The peak at the lower molecular weight represents free nucleotides. Both ADP and ATPγS-loaded DFCP1 K193A elute primarily as a single peak, corresponding to the molecular weight of a DFCP1 monomer. ADP-loaded DFCP1 T189A elutes as a complete monomer, and no side peaks corresponding to higher molecular weights are detectable.

Structure prediction using AlphaFold ^11^, as well as structural homology modelling using the Phyre2 server ^12^ revealed that the DFCP1 P-Loop domain shows structural homology to large nucleotide binding proteins, especially to the GTPases Atlastin and GBP1 (Fig.1b,c, Extended Data Fig.1c). Based on this finding, we tested if DFCP1 is able to bind nucleotides. We purified the N-terminus of DFCP1, including the predicted nucleotide-binding domain (Extended Data Fig.2a). To measure nucleotide binding, we used N-methylanthraniloyl (mant)-modified nucleotides, mantATP and mantGTP, which are not fluorescent in solution but emit fluorescence upon binding to a protein. We observed no binding to GTP within the time course of our measurements, whereas Cdc42 – a GTP-binding protein – showed robust binding to GTP (Extended Data Fig.2b). In contrast, DFCP1 efficiently and specifically bound to ATP (Fig.1d). DFCP1 also bound to ADP (Extended Data Fig.2c), suggesting that it could act as a molecular switch, depending on the loaded nucleotide. Binding of ATP/ADP to DFCP1 was rapid, with the reaction reaching saturation within minutes (Fig.1d). To address whether DFCP1 has ATPase activity, we measured DFCP1-dependent phosphate release from ATP. Indeed, adding ATP to the purified DFCP1 ATPase domain resulted in the release of free phosphate (Fig.1e). Thus, DFCP1 is a functional ATPase.

To understand the functional importance of the ATP binding and hydrolysis by DFCP1, we aimed to generate mutations that affect these biochemical properties. We used a Phyre2-generated homology model based on Atlastin, which showed the highest degree of sequence similarity (Fig.1b,c). Using this model, we performed structural alignments against other nucleotide-binding proteins (Atlastin, GBP1, N-RAS and K-RAS) to identify key residues necessary for nucleotide binding and hydrolysis.

NTPases have a characteristic GKS motif, which is critical for nucleotide binding by coordinating a magnesium ion ^13^. We identified this motif at residues 192-194 of DFCP1. Based on homologies with the small GTPases K-RAS and N-RAS, we further identified the residues T189 and G190 as potentially critical residues. T189 is at the same position as the G12 residue of K-RAS, whereas G190 corresponds to G13 in K-RAS. Both amino acids are critical for K-RAS GTPase activity (Fig.1b,c).

Based on these predictions, we generated and purified DFCP1 nucleotide-binding defective mutants (Extended Data Fig.2a). Similar to other nucleotide binding proteins, mutation of the GKS motif (K193A, S194N) resulted in a loss of ATP binding (Fig.1d). The same was the case with mutations of the K-Ras G12/G13 analogous amino acids (T189V and G190V) (Fig.1d).

We also aimed to identify an ATPase-defective DFCP1 mutant and found that the mutation T189V, which corresponds to the hydrolysis-defective K-Ras G12V allele, showed slightly reduced hydrolysis activity, but showed also reduced nucleotide binding (Fig.1d,e). Importantly, however, another mutation of T189, T189A, showed robust nucleotide binding but only minimal ATP hydrolysis activity (Fig.1f,g), thus constituting an ATP-locked DFCP1 allele.

### DFCP1 dimerizes upon ATP binding

Many NTPases, such as Dynamin and Atlastin, oligomerize as a consequence of nucleotide binding ^14^. To elucidate if DFCP1 shows nucleotide-dependent oligomerization, we loaded the purified DFCP1 N-terminus with ADP or the non-hydrolysable ATP analogue, ATPγS, and performed size exclusion chromatography. Interestingly, whereas ADP-bound DFCP1 migrated as a monomer, ATPγS-loaded DFCP1 migrated as dimer (Fig.1h, Extended Data Fig.2d). As a control, we performed the same assay using DFCP1 K193A, the nucleotide-binding defective mutant we identified. This mutant migrated as a monomer and did not dimerize in the presence of ATPγS (Fig.1h, Extended Data Fig.2d), in line with our findings that dimerization of DFCP1 is ATP-dependent. Surprisingly, DFCP1 T189A, which binds ATP but is hydrolysis-deficient, also migrated as a monomer in the presence of ATPγS (Fig.1h, Extended Data Fig.2d). This suggests that dimerization of DFCP1 could be required for ATP hydrolysis.

### DFCP1 ATP binding and hydrolysis are necessary for efficient omegasome constriction

To address whether cells expressing the DFCP1 mutants show defects in omegasome formation, we generated a knockout (KO)-based complementation system (Extended Data Fig.3a-d). DFCP1 KO cells were transduced with lentiviral vectors expressing low levels of either mNeonGreen (mNG)-tagged DFCP1 wild-type (WT) or the two ATP binding or hydrolysis mutants K193A or T189A, in addition to mCherry-(mCh)-p62 as an autophagosomal marker. Structured illumination microscopy (SIM) revealed that mNG-DFCP1 WT localized to ring shaped omegasomes containing LC3B and p62 (Fig.2a). To assess omegasome dynamics, the cells were starved with EBSS for 15 min and images were acquired for 10 minutes with one frame taken every 2 s in starvation conditions (Fig.2b, movie 1).

**Fig.2:**
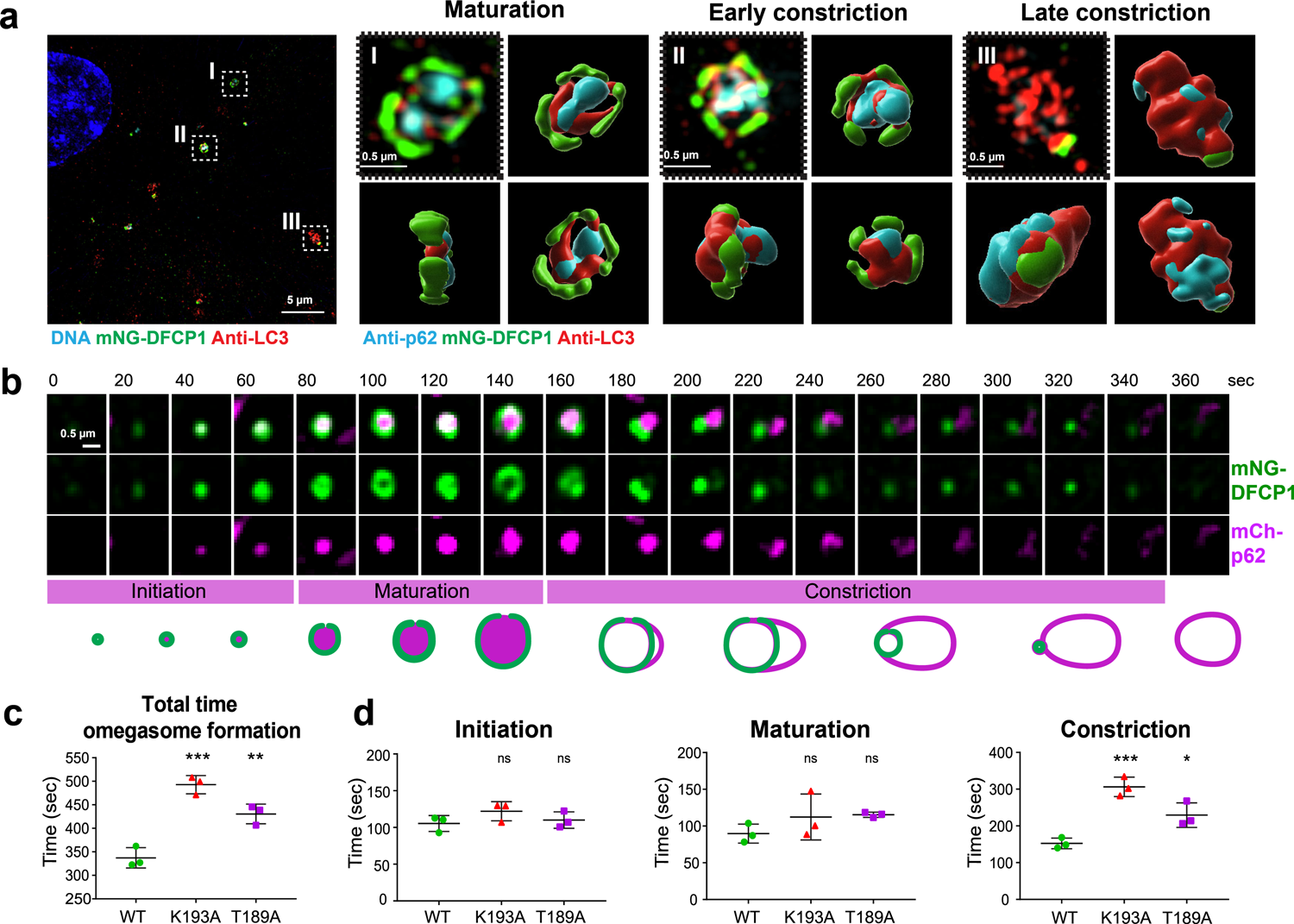
DFCP1 mutants show delayed omegasome constriction. Fixed or live imaging of U2OS DFCP1 KO lines stably expressing mNG-DFCP1 WT, K193A or T189A as indicated: (a) 3D reconstruction of omegasomes in maturation and constriction phases. U2OS DFCP1 KO lines stably expressing mNG-DFCP1 WT were starved in EBSS, fixed, stained for p62, LC3B and mNG and SIM imaging was performed. Shown are examples from three different stages of omegasome formation, and their reconstruction from different angles. Scale bar: 5 µm, insets: 0.5 µm. (b) U2OS DFCP1 KO lines stably expressing mNG-DFCP1 WT and mCh-p62. Cells were starved in EBSS and imaged live to capture omegasome formation. Omegasome formation was divided into three phases as indicated. The scheme illustrates how DFCP1 and p62 co-localize throughout an omegasomes’ life cycle. Quantification of events in (b) and (c). Scale bar: 0.5 µm. (c) Quantification of the duration of an omegasome life cycle, with a DFCP1 punctum as beginning and end point. Each plotted point represents the mean value of one experiment. Bars: mean ± SD, 3 experiments, analysing in total 58 (WT), 16 (K193A) and 49 (T189A) omegasomes. One-way Anova with Dunnett’s post-hoc test, comparing against WT. (d) Duration of the individual phases of omegasome formation. Each plotted point represents the mean value of one experiment. Bars: mean ± SD, 3 experiments. Statistics: Ordinary One-way Anova followed by Dunnett’s multiple comparisons test, comparing against WT. 3 experiments, analyzed omegasomes: Initiation: WT=76, K193A= 54, T189A=81; Maturation: WT=99, K193A=81, T189A=117; Constriction: WT=78, K193A=47, T189A=99.

Autophagosome formation at omegasomes is a reproducible course of events ^4, 15^. Based on published data and the tracking of DFCP1 WT omegasomes by live microscopy, we divided the omegasome formation process into three phases – initiation, maturation, and constriction (Fig.2b). The initiation phase is characterized by a DFCP1 spot which is formed de novo, and to which p62 and LC3B are recruited a few seconds later ^15, 16^ Fig.2b, Extended Data Fig.4d). The spot grows and forms a ring-like structure, as previously observed ^4 15^. We have defined the appearance of a ring with a visible lumen as the start for the maturation phase. LC3B and p62 are recruited to the initial DFCP1 spot and during the maturation, the DFCP1 ring grows, and LC3B and p62 span the lumen forming a disk-like structure (Fig.2a; Extended Data Fig.4b,e ^15–17^). At the end of maturation, a p62 and LC3B-labelled phagophore extrudes out of the ring and forms a pocket (Fig.2a; Extended Data Fig.4b,e). This separation of the DFCP1-positive omegasome from the p62/LC3B positive phagophore initiates the third phase – the constriction of the omegasome. The phagophore bends, buds out and is separated from the DFCP1-positive omegasome, thereby forming the nascent autophagosome. During this process, the shrinking omegasome remains connected to the growing autophagosome, which is positive for both p62 and LC3B, but also weakly for DFCP1 (Fig.2a; Extended Data Fig.4b,e). Finally, the weak DFCP1 signal leaves the autophagosome, and DFCP1 at the collapsed omegasome disappears, leaving a p62/LC3B-positive-finished autophagosome (Fig.2a).

We next analysed the dynamic formation of omegasomes in DFCP1 mutant cells. Whereas the omegasomes appeared morphologically similar to the WT omegasomes (Extended Data Fig.4b,e), detailed analysis and tracking revealed a significant delay in the omegasome formation process (Fig.2c, Extended Data Fig.4a, movie2). While DFCP1 WT used approximately 330 sec to form omegasomes as previously described ^16^ ^4, 15^, the ATP-binding mutant DFCP1 K193A used 500 sec, whereas the ATPase-defective mutant DFCP1 T189A needed 450 sec to complete the process.

To understand which step in omegasome biogenesis was affected in the mutants, we measured the duration of the three individual phases. Omegasomes in cells expressing DFCP1 WT used approximately 100 sec for each of the two first phases, whereas the last phase took 150 sec. Surprisingly, cells expressing either of the two DFCP1 mutants showed a nearly unchanged duration for the first two phases. In contrast, the constriction phase was markedly delayed compared to WT (Fig.2d, Extended Data Fig.4c). We confirmed these findings also using DFCP1 knockdown cell lines, which were rescued with stable expression of low levels of siRNA resistant WT or mutant mNG-DFCP1 in combination with SNAP-LC3B (Extended Data Fig.3e-g, Extended Data Fig.4d-i, movie 3, movie 4). Based on these data, we conclude that the ATP binding and hydrolysis activities of DFCP1 are necessary for the efficient constriction of omegasomes.

### DFCP1 ATPase mutants cause increased numbers of omegasomes

Based on our finding that DFCP1 ATPase is required for omegasome constriction, we asked which consequences it has for autophagy. Upon starvation with EBSS, cells expressing the ATP-binding or ATPase-defective mutants showed an increased number of DFCP1 puncta compared to DFCP1 WT, and both mutants had a higher number of larger omegasomes (Fig.3a, b).

**Fig.3:**
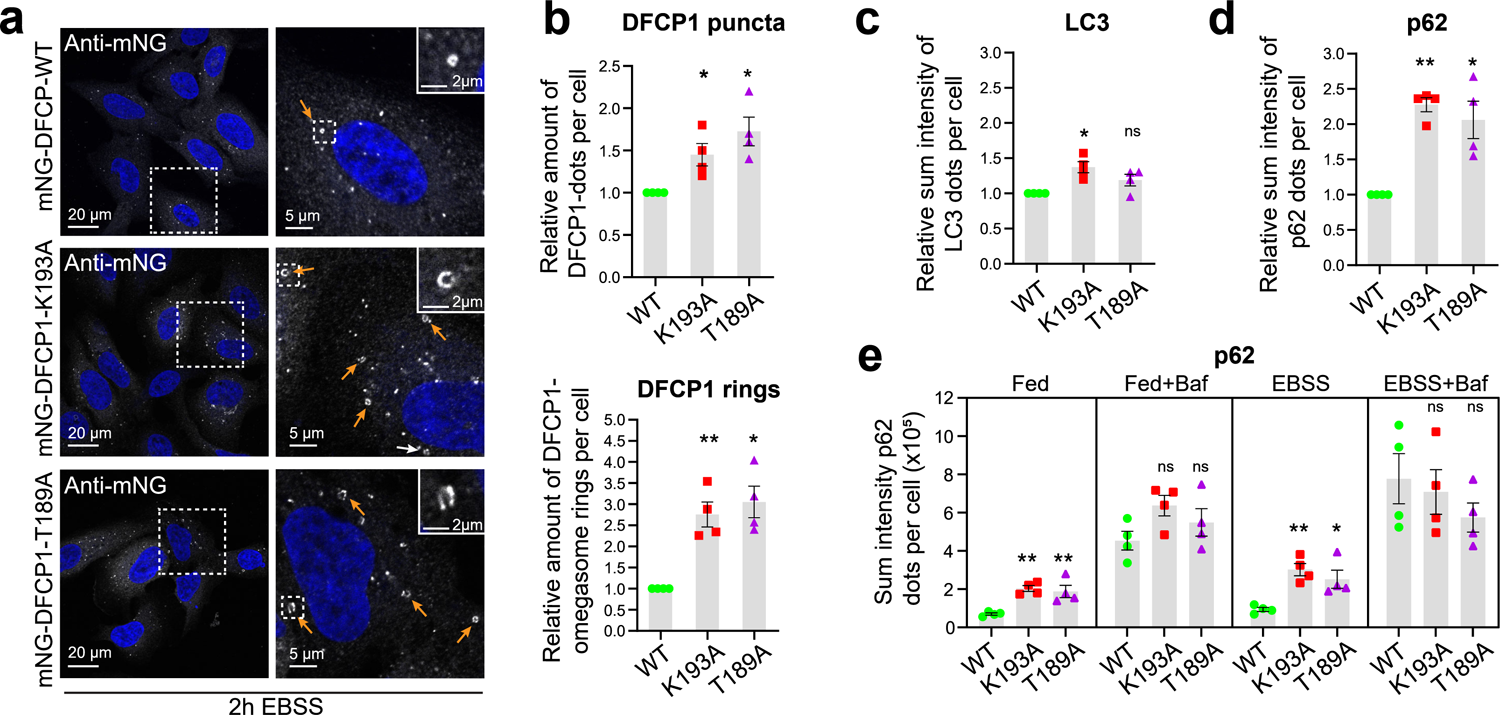
p62 accumulates in DFCP1 mutants defective in ATP binding or hydrolysis. Fixed imaging of U2OS DFCP1 knock-down rescue lines stably expressing mNG-DFCP1 WT, K193A or T189A as indicated: (a) Confocal images showing the accumulation of different omegasome stages in DFCP1 WT and mutant cells. U2OS cells stably expressing siRNA resistant mNG-DFCP1 WT, K193A or T189A, were depleted of endogenous DFCP1 by siRNA transfection for 2 d in complete medium before starvation for 2 h in EBSS. The cells were fixed, stained with anti-mNG antibody, and analyzed by confocal microscopy. Arrows indicate DFCP1 positive omegasome rings. Scale bar: 20 µm, inset: 5 µm or 2 µm. Representative image for quantifications in (b). (b) Quantification of the relative amount of mNG-DFCP1 dots and rings per cell. Error bars denote mean ± SEM, 4 experiments. Each plotted point represents the mean value of one experiment. One-sample t-test: * p < 0.05; ** p < 0.01. Analyzed cells per condition: WT: 643, K193A: 650, T189A: 632. (c) Cells treated as in (a). The cells were fixed, stained with anti-LC3B and analyzed by confocal microscopy. Relative sum intensity of LC3B dots per cell. Error bars denote mean ± SEM, 4 experiments. Each plotted point represents the mean value of one experiment. One-sample t-test: * p < 0.05; ns: not significant. Analyzed cells per condition: WT= 458, K193A= 461, T189A= 446. (d) Cells treated as in (a). The cells were fixed, stained with anti-p62 and analyzed by confocal microscopy. Relative sum intensity of p62 dots per cell. Error bars denote mean ± SEM, 4 experiments. Each plotted point represents the mean value of one experiment. One-sample t-test: * p < 0.05; ** p < 0.01. Analyzed cells per condition: WT= 643, K193A= 650, T189A= 632. Same dataset as in b. (e) U2OS cells stably expressing siRNA resistant mNG-DFCP1 WT, K193A or T189A, were depleted of endogenous DFCP1 by siRNA transfection for two days. Shown are fed cells (complete medium), EBSS-starved cells (EBSS for 2h), in the presence or absence of 100 nM Bafilomycin A1 to assess autophagic flux. Cells were fixed, stained with anti-p62 antibody, and analyzed by confocal microscopy. Graphs represent quantification of the sum intensity of p62 dots per cell in the different conditions as indicated. Error bars denote mean ± SEM, 4 experiments. Each plotted point represents the mean value of one experiment. One-way Anova, Dunnett’s multiple comparisons test comparing against WT within each treatment group. * p < 0.05; ** p < 0.01; ns: not significant. In total >400 cells were analyzed per cell line per condition.

The increased number of omegasomes in the DFCP1 mutants can be either a consequence of increased omegasome formation or prolonged persistence. To investigate whether omegasome formation was increased, we made use of another omegasome marker, WIPI2 ^9, 10^. WIPI2 is recruited by PtdIns3P to phagophores originating at omegasomes and appears nearly at the same time as DFCP1 but dissociates earlier ^9^. Thus, WIPI2 represents a good marker for omegasome formation independently of DFCP1. We analysed endogenous WIPI2 puncta 2 hrs after EBSS treatment in DFCP1 WT and mutants. As expected, a portion of the WIPI2 dots co-localized with DFCP1 positive omegasomes, likely representing early stages of autophagosome formation (Extended Data Fig.5a). Interestingly, although the number of DFCP1 puncta was higher in cells expressing the mutants, neither the number of WIPI2 puncta nor the portion of DFCP1 positive WIPI2 puncta was changed (Extended Data Fig.5b), suggesting that omegasome formation was not increased. On the other hand, the portion of DFCP1 puncta positive for WIP2 was reduced in cells expressing the mutants, indicating that the DFCP1 positive omegasomes that accumulated in the mutant cells likely represent a later, WIPI2 negative, stage of omegasomes. We conclude that whereas the number of omegasomes increases in the DFCP1 mutant cells, the onset of omegasome formation is not affected.

### Autophagic flux of p62 is compromised in DFCP1 ATPase mutants

It has remained a paradox that depletion of DFCP1 by siRNA does not have any effect on the flux of LC3B (Axe 2008). However, how LC3B flux is affected upon specific modulate of DFCP1 ATPase activity was not clear. First, we measured the sum intensity of endogenous LC3B puncta in DFCP1 WT and DFCP1 mutant rescue cells by automated analysis of confocal micrographs. As expected, numerous LC3B puncta were detected upon EBSS starvation in DFCP1 WT cells (Extended Data Fig.5c). Interestingly, DFCP1 ATP-binding deficient K193A mutant cells showed a small, but significant increase in the sum intensity of LC3B dots, whereas cells expressing the ATP-locked T189A mutant only showed a tendency to accumulate LC3B as compared to DFCP1 WT (Fig.3c, Extended Data Fig.5c).

It has been reported that p62 localizes to phagophores which are formed at omegasomes, and that this recruitment occurs independently of LC3B ^18^. In line with this, we noticed that while LC3B was present on many other structures in addition to omegasomes, the majority of p62 puncta localized with DFCP1 positive omegasomes (Extended Data Fig.5c). Importantly, when we measured the sum intensity of endogenous p62 puncta, both DFCP1 ATPase mutant cell lines showed a more than two-fold increase in p62 dot intensity as compared to DFCP1 WT cells upon EBSS treatment (Fig.3d). Moreover, in cells expressing high amounts of DFCP1, p62 hyperaccumulated inside DFCP1 ATPase deficient omegasomes (Extended Data Fig.5d).

To address whether the increased level of p62 dots observed in the mutants could be explained by an impaired autophagic flux, we measured the sum intensity of p62 dots per cell in fed or EBSS starved cells in the presence or absence of Bafilomycin A1, which inhibits lysosomal degradation activity. This analysis confirmed our previous result, with a close to three-fold increase in p62 dot sum intensity in the EBSS treated mutant cells (Fig.3e). In addition, we observed a similar effect in fed cells, indicating that also basal autophagy was impaired in DFCP1 depleted or ATPase deficient cells (Fig.3e, Extended Data Fig.6a-e). Addition of Bafilomycin A1 indeed prevented the lysosomal degradation of p62, as the sum intensity of p62 dots clearly increased (Fig.3e). Importantly, when lysosomal degradation was blocked with Bafilomycin A1, there was no difference in the p62 levels between DFCP1 WT or ATPase defective cells (Fig.3e). This indicates that p62 accumulates in the DFCP1 ATP-binding and ATPase defective mutants due to a delayed autophagic flux, rather than increased onset of autophagy, consistent with our finding that the number of WIPI2 puncta were unaffected. Taken together, our data show that DFCP1 is required for efficient autophagic flux of p62, and that this depends on its ability to bind and hydrolyse ATP.

### DFCP1 ATPase is required for selective autophagy

As expected from our and published ^4^ results on LC3B lipidation upon DFCP1 depletion, we found that DFCP1 is not involved in bulk autophagy, using cytoplasmic mKeima reporter assays ^19, 20^ in U2OS and RPE-1 cells (Extended Data Fig.6f-i). However, since we observed that p62 accumulates in DFCP1 ATPase mutants in both fed and starved conditions, we asked if the DFCP1 ATPase mutants are impaired in selective autophagy.

p62 has been implicated in several types of selective autophagy, such as aggrephagy and mitophagy ^21–23^. To induce aggrephagy, we treated stably expressing mCh-DFCP1 cells with Puromycin. Puromycin treatment leads to the formation of large p62-containing aggregates, which are heavily ubiquitinated and cleared by autophagy ^24^. Notably, DFCP1 localized to a subset of these aggregates, forming characteristic omegasomes positive for LC3B (Fig.4a, Extended Data Fig.7a,b). Strikingly, DFCP1 KO cells had more than a two-fold increase in p62 aggregates upon Puromycin treatment compared to the WT (Fig.4b). This could be rescued by WT DFCP1, but not the ATPase mutants (Fig.4c, Extended Data Fig.7c). Thus, DFCP1 and its ATPase activity are necessary for efficient aggrephagy.

**Fig.4:**
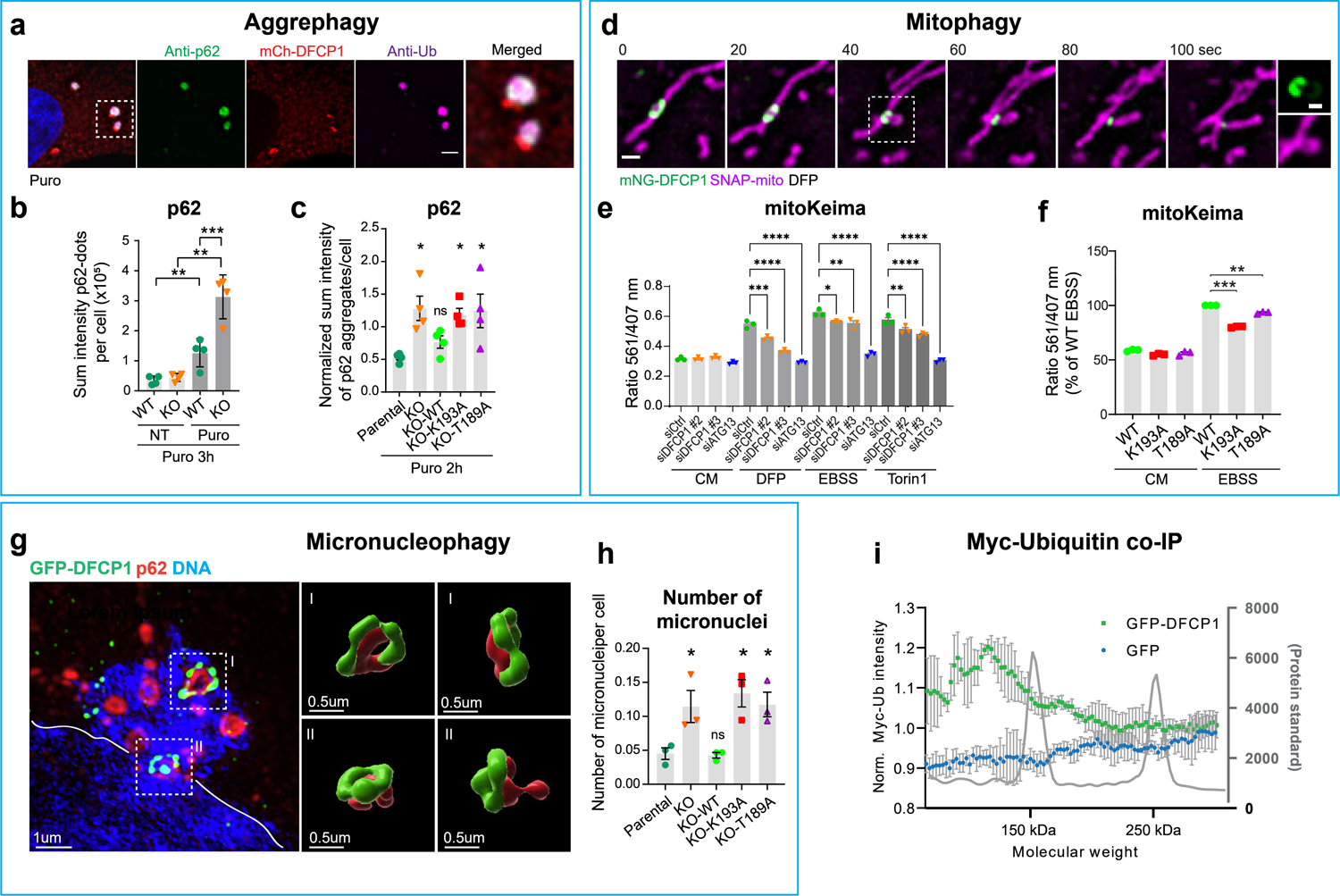
DFCP1 ATPase regulates selective types of autophagy and binds ubiquitin. (a) U2OS cells stably expressing mCh-DFCP1 were treated with 10 µg/ml Puromycin for 3 h, fixed, and stained with antibodies against p62 and ubiquitin. Representative image for >15 cells, scale bar: 2 µm. (b) U2OS WT and DFCP1 KO cells were treated or not with 2.5 µg/ml µg Puromycin for 3 h, fixed, stained with anti-p62 antibody, and analyzed by confocal microscopy. Graphs represent quantification of the relative sum intensity of p62 dots per cell. Statistics: unpaired t-test, 4 experiments. Each plotted point represents the mean value of one experiment. Cells analyzed per condition: WT-NT: 1175, KO-NT: 1042, WT-Puro: 1465, KO-Puro: 1381. (c) Parental U2OS, DFCP1 KO and KO rescue lines stably expressing mNG-DFCP1 WT, K193A or T189A were treated with 2.5 µg/ml Puromycin for 2 h, fixed, stained with anti-p62 antibody, and analyzed by confocal microscopy. Graphs represent quantification of the relative sum intensity of p62 aggregates per cell. Values have been normalized by the mean of the mean. Bars: mean ± SEM, 4 experiments. Statistics: One-way Anova with Dunnett’s posthoc test, comparing against U2OS parental. Cells analyzed per condition: parental: 1732, KO:1716, KO-WT: 2392, KO-T189A: 2601, KO-K193A: 2263. (d) Example of a mitophagy event. U2OS cells stably expressing mNG-DFCP1 WT and SNAP-mito were treated for 18 h with 0.5 mM DFP and imaged live. The inset shows a piecemeal of mitochondria, which is engulfed by the phagophore. Note that the phagophore is weakly positive for mNG-DFCP1 (inset). Representative movie for > 10 events. Scale bar: 1 µm, inset 0.5 µM. (e) RPE-1 cells induced to express a mitochondrial mKeima probe were depleted for DFCP1 for 2 d and were cultured for 16 h in either DFP, EBSS or Torin1, harvested and analyzed by Flow Cytometry to determine the ratio of lysosome-localized mito-mKeima to free, cytoplasmic mito-mKeima. Bars: ± SEM, 3 experiments. Statistics: One-way Anova followed by Dunnett’s multiple comparisons test. (f) RPE-1 cells stably expressing mitochondrial mKeima probe and siRNA resistant mNG-DFCP1-WT, K193A, or T189A were depleted of endogenous DFCP1 by siRNA transfection for 2d in complete medium (CM) before starvation for 16 h in EBSS to induce mitophagy. Cells were harvested and analyzed by Flow Cytometry to determine the ratio of lysosome-localized mito-mKeima to free, cytoplasmic mito-mKeima. Values were normalized to EBSS-treated WT. Bars: ± SEM, 3 experiments. Statistics: One-sample t-test. (g) GFP-DFCP1 knockin cells have been fixed and stained with p62 and GFP-antibodies and Hoechst33342 and analyzed by SIM-microscopy. Shown is a micronucleus decorated with omegasome rings, which have been 3D-rendered and are shown from different angles. The border of the main nucleus is indicated by a white line. Scale bar: 1 µm, inset: 0.5 µm. (h) U2OS parental cells, DFCP1-KO and KO rescue lines stably expressing mNG-DFCP1 WT, K193A or T189A were grown in complete medium, fixed, stained with Hoechst33342, analyzed by confocal microscopy and the number of micronuclei determined. Error bars denote mean ± SEM, 3 experiments. Each plotted point represents the mean value of one experiment. One-way ANOVA, Dunnett’s post hoc test: * p < 0.05; ns: not significant. Cells analyzed per condition: parental=788; KO= 703; WT= 732, K193A= 779, T189A= 766. Same data set used to quantify p62 intensity in Extended Data Fig.6b, e. Images of MN in Extended Data Fig.7h. (i) Quantification of co-IP of transfected HeLa cells from WB membranes as shown in Extended Fig.8b. Line plot of Myc-Ub intensities quantified from the IP samples of WB membranes in the region indicated by the protein standard. Data is normalized to the mean experiment intensity. n=3 experiments +/- SD.

To investigate if DFCP1 is necessary for autophagic degradation of mitochondria, we used the iron chelator Deferiprone (DFP) to induce mitophagy ^25^ in cells stably expressing mNG-DFCP1 and a mitochondrial marker (mitoSNAP) and performed live imaging. We observed that DFCP1 formed omegasomes at the surface of mitochondria, and that mitochondrial fragments were channelled through the omegasome ring (Fig4.d, Extended Data Fig.7d, movie 5, movie 6). Importantly, the mitochondrial fragments were surrounded by the weakly DFCP1-positive phagophore, indicating that it had been engulfed by the autophagosome (insets Fig.4d, Extended Data Fig.7d). Following engulfment and constriction of the omegasome, autophagosomes were released from the mitochondria (Extended Data Fig.7d).

The importance of DFCP1 for mitophagy was quantified using RPE-1 cells stably expressing a mitochondrial mKeima probe, mito-mKeima ^19, 26^. When mito-mKeima is transported to acidic lysosomes for degradation, the excitation maximum changes and mitophagy can be determined by flow cytometry as an increased ratio of lysosomal mito-mKeima. While control cells displayed a nearly two-fold increase in lysosomal mito-mKeima when mitophagy was induced, DFCP1 depleted cells had a strongly reduced capacity to deliver mito-mKeima to lysosomes (Fig.4e, Extended Data Fig.7e).

To address whether DFCP1 ATPase activity is necessary for mitophagy, we stably expressed DFCP1 WT or mutants in the mito-mKeima RPE-1 cells depleted of endogenous DFCP1 and performed mitophagy assays (Fig.4f, Extended Data Fig.7f,g). Both ATP-binding and ATPase defective mutants of DFCP1 were impaired in mitophagy (Fig.4f, Extended Data Fig.7f,g), indicating that the ATPase activity of DFCP1 is necessary for efficient mitophagy.

When examining the phenotype of U2OS DFCP1 KO cells, we noticed that they contained an increased number of micronuclei. Image quantifications showed that DFCP1 KO and ATPase deficient cells had a more than two-fold increase in the number of micronuclei per cell as compared to parental or DFCP1 WT cells (Fig.4h, Extended Data Fig.7h). This phenotype was also confirmed in two independent clones of A431 DFCP1 KO cells (Extended Data Fig.7k,l). Similarly, U2OS cells acutely depleted for DFCP1 showed an increase in the number of micronuclei per cell (Extended Data Fig.7i,j). Importantly, we noticed that DFCP1 localized to a subset of micronuclei, colocalizing with p62. Super-resolution microscopy revealed that endogenously GFP-tagged DFCP1 formed characteristic omegasome rings at the surface of micronuclei together with p62 (Fig.4g). These results suggest that DFCP1 promotes autophagic degradation of micronuclei, in addition to mitochondria and protein aggregates.

### DFCP1 associates with ubiquitinated proteins

How could it be explained mechanistically that DFCP1 mediates selective autophagy whereas it is dispensable for bulk autophagy? We noticed that omegasomes stain positive with antibodies against conjugated ubiquitin (Extended Data Fig.8a, c). As in the case of aggrephagy (Fig.4a), also omegasomes decorating mitochondria or micronuclei were positive for ubiquitin (Extended Data Fig.8a, c), raising the possibility that DFCP1 could interact with ubiquitinated cargoes. To investigate this notion, we co-transfected HeLa cells with Myc-epitope-tagged ubiquitin and GFP-DFCP1 and investigated whether affinity isolated GFP-DFCP1 would co-precipitate ubiquitinated proteins. Interestingly, whereas no ubiquitinated proteins co-precipitated with GFP as revealed by immunoblotting with anti-Myc, a smear of Myc-Ubiquitin positive bands was pulled down with GFP-DFCP1 (Fig.4i, Extended Data Fig.8b). This indicates that DFCP1 associates with ubiquitinated cargoes and provides a plausible model for its function.

## Discussion

DFCP1 translocates to omegasomes in a PtdIns3P dependent manner following amino acid starvation ^4^. Although DFCP1 has been widely used as an omegasome marker, its function was completely unknown. Here, we have found that DFCP1 associates with ubiquitinated cargoes and uses its ATPase activity to mediate a timely constriction of omegasomes in selective autophagy.

Our bioinformatic analyses showed that DFCP1 bears highest similarity to large GTPases such as GBP1 and Atlastins (Fig.1b), and we were surprised to find that it functions as an ATPase. A similarly altered nucleotide specificity has been found for EHD-related proteins, which are related to Dynamin, but are selective for ATP instead of GTP ^27^.

Threonine-189, which we found essential for ATP hydrolysis, is conserved in the GTPase Irga6. In Irga6, residues in switch I are able to contact the ribose of the nucleotide bound in the opposing protein, which stabilizes dimerization^28^. This dimerization promotes GTP hydrolysis. Importantly, a mutation in Threonine-78 of Irga6 ^29^, which corresponds to Threonine-189 in DFCP1, interferes with dimerization and strongly reduces nucleotide hydrolysis. We found that wild-type but not ATPase-defective (T189A) DFCP1 forms an ATP-dependent dimer. This suggests that dimerization of DFCP1 is required for effective nucleotide hydrolysis and that DFCP1 employs a similar mechanism as Irga6 for nucleotide hydrolysis.

Our functional data indicate that the DFCP1 ATPase is involved in the maturation of omegasomes, specifically during omegasome constriction. The inhibited constriction in DFCP1 mutants bears a striking similarity to the constriction of membrane necks by Dynamin and the phenotypes of both hydrolysis-defective and nucleotide binding Dynamin mutants. Dynamin binds to membranes by its C-terminal PH domain and forms spiral-like assemblies at membrane necks. Cycles of nucleotide loading and hydrolysis cause these assemblies to constrict, which ultimately drives membrane scission ^30^. Similar to Dynamin, also DFCP1 can bind to membranes by its two C-terminal FYVE domain, which contribute to anchor the protein to the membrane. It is tempting to speculate that the nucleotide hydrolysis cycle of DFCP1 could lead to repeated dimerization and dissociation of the ATP-binding domain, which would lead to a stepwise constriction of the omegasome (Extended Data Fig.9).

Since p62 positive forming autophagosomes were connected to the DFCP1 rings for the whole duration of their constriction period, we speculate that ring constriction is likely an important prerequisite for sealing of the extended phagophore, and to release the nascent autophagosome (Extended Data Fig.9). Both the ATP-binding defective mutant and the ATP-locked mutant showed a similar phenotype, which can probably be attributed to our finding that both mutants are unable to dimerize. Physiologically, this slowed-down constriction of the omegasome translates to a delay in autophagic flux of p62.

Previously it had been concluded that depletion of DFCP1 has no effect on autophagic flux, measured by LC3 following amino acid starvation ^4^. When we specifically measured bulk autophagy using a free cytoplasmic Keima probe, we confirmed that DFCP1 does not play a role in bulk autophagy. In contrast, we found a striking dependence on DFCP1 in selective autophagy. This was unanticipated given the fact that DFCP1-containing omegasomes are typically induced by amino acid starvation. However, such starvation does not only induce bulk autophagy but also accumulation of various ubiquitinated objects ^25^ ^31^, and this is probably the explanation why omegasomes are observed after starvation.

In addition to p62-dependent aggrephagy, we also found that efficient degradation of mitochondria and micronuclei depends on the ability of DFCP1 to bind and hydrolyse ATP. Further studies are needed to establish the exact functions of DFCP1 in mitophagy and micronucleophagy, but it is striking that several types of selective autophagy depend on DFCP1. While it is interesting that KO of DFCP1 results in accumulation of micronuclei, we cannot exclude the possibility that this increase is a secondary effect on cell division caused by impaired selective autophagy ^32, 33^. However, the presence of multiple DFCP1-labelled omegasomes on micronuclei supports a direct role for DFCP1 in micronucleophagy.

In the three types of selective autophagy we studied, we observed both ubiquitinated proteins and DFCP1 around the cargoes. Together with our finding that DFCP1 co-precipitates with ubiquitinated proteins and a recent study which identified ubiquitin as the top interactor of DFCP1 ^34^, this suggests that DFCP1 serves to sequester ubiquitinated cargoes into omegasomes. This bears resemblance to another PtdIns3P-binding protein, HRS, which sequesters ubiquitinated cargoes into intraluminal vesicles of endosomes ^35^. In addition to binding ubiquitinated cargoes, HRS becomes ubiquitinated itself ^36^, and our data do not exclude the possibility that DFCP1 is also ubiquitinated.

In conclusion, we have identified an evolutionarily conserved Dynamin-related ATPase module that controls the early phase of selective autophagy (Extended Data Fig.9). It is interesting to note that biogenesis of the replication compartments of certain viruses, including SARS-CoV-2, depends on DFCP1 ^37^, and our data thus imply that constriction of ER-derived membranes is a critical step in SARS-CoV-2 replication. DFCP1 recognizes ubiquitinated cargoes which are included in omegasomes, and ATP binding and hydrolysis of DFCP1 trigger omegasome constriction and thereby the biogenesis of cargo-containing autophagosomes. In future studies it will be interesting to reveal the interplay between DFCP1 and cargoes, and to understand the structural biology of the novel ATPase

## Methods

### Antibodies

| Primary antibodies |  |  |  |  |
| --- | --- | --- | --- | --- |
| Target | Species | Source | Identifier | Concentration |
| DFCP1 (E9R6P) | rabbit | Cell Signaling Technology | mAb #85156 | WB 1:500 |
| mNeonGreen [32F6] | mouse | Chromotek | 32f6-100 | IF 1:500 |
| GFP | mouse | Roche | 11814460001 | WB 1:500 |
| Myc | rabbit | abcam | ab9106 | WB 1:20 000 |
| beta-actin | mouse | Sigma-Aldrich | A5316 | WB 1:20 000 |
| LC3B | rabbit | MBL | PM036 | IF 1:500 |
| LC3B | rabbit | Cell Signaling Technology | 2775s | WB 1:1000, IF 1:100 |
| WIPI2 | mouse | abcam | ab105459 | IF 1:200 |
| p62 | rabbit | MBL | PM045 | WB 1:10 000, IF 1:500 |
| p62 | guinea pig | Progen | GP62-C | IF 1:250 |
| Ubiquitin (FK2) | mouse | EMD Millipore Corp. | 04-263 | IF 1:400 |
| Ubiquitin-K63 | rabbit | Millipore | 05-1308 | IF 1:200 |
| Tom20 | mouse | BD | 612278 | IF 1:100 |
| Atg13 | rabbit | Cell Signaling Technology | #13468 | WB 1:1000 |
| Secondary antibodies |  |  |  |  |
| Alexa488-anti-mouse | donkey | Jackson | 715-545-151 | 1:500 |
| Alexa488-anti-rabbit | donkey | Jackson | 711-545-152 | 1:500 |
| Alexa568-anti-mouse | donkey | Molecular Probes | A10037 | 1:500 |
| Alexa568-anti-rabbit | donkey | Molecular Probes | A10042 | 1:500 |
| Alexa647-anti-mouse | donkey | Jackson | 715-605-150 | 1:500 |
| Alexa647-anti-rabbit | donkey | Jackson | 711- 605 -152 | 1:500 |
| Alexa647-anti-guinea pig | donkey | Jackson | 706-605-148 | 1:500 |

### Constructs

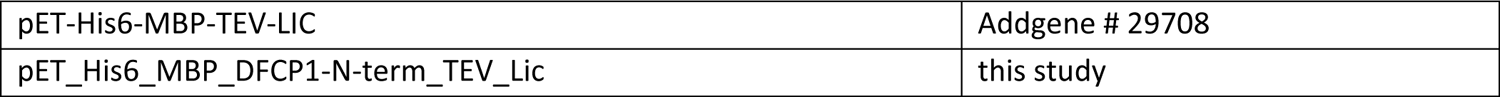
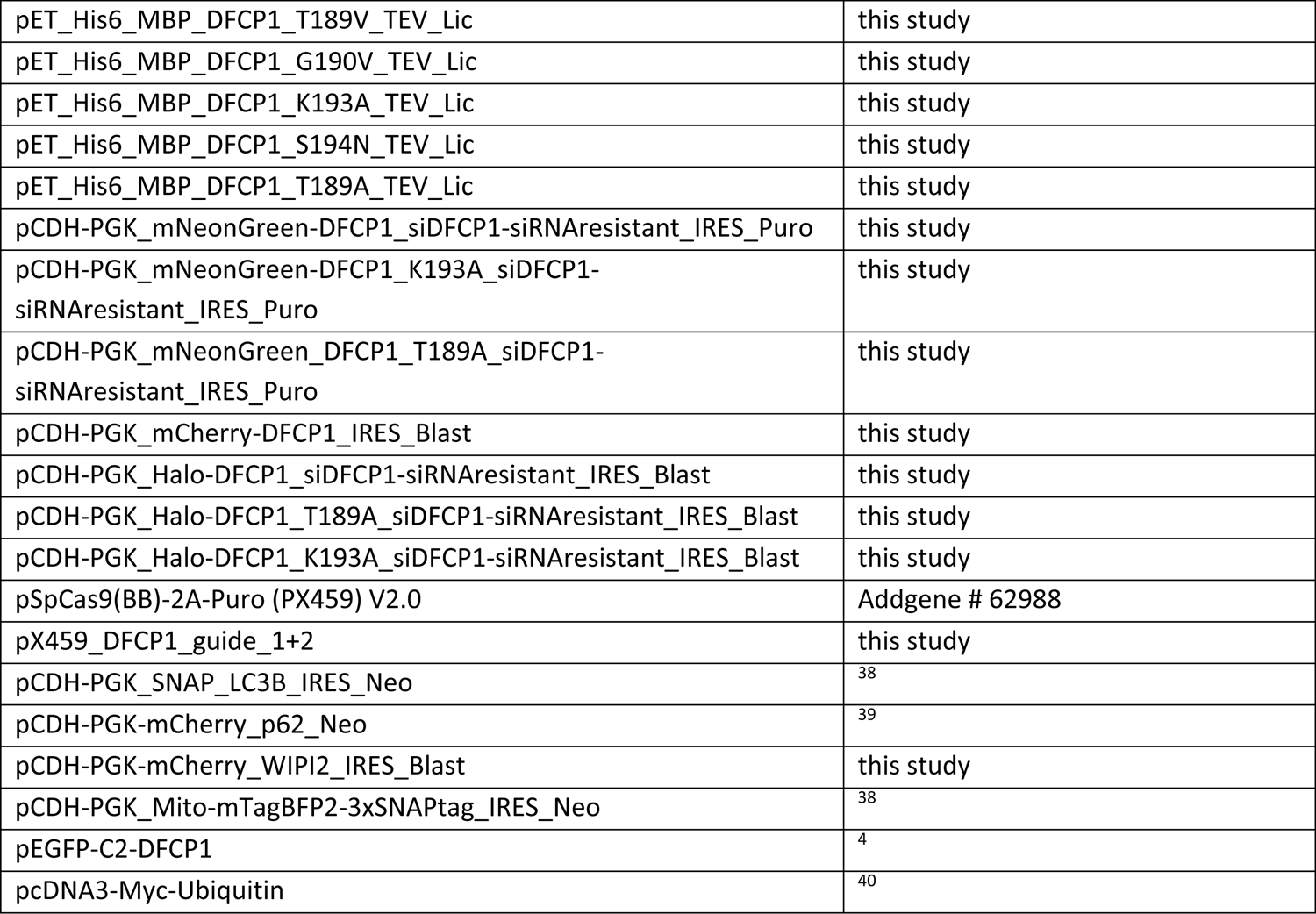

### Structure prediction, homology modelling and structural analysis

We performed structure prediction using AlphaFold ^11^. Homology modelling was also performed using the Phyre2 server, using DFCP1 amino acids 1-410 as target sequence. The complete output of the Phyre2 modelling and from AlphaFold is attached as Supplementary Information. For structural comparison with Phyre2, we chose the prediction model with the highest confidence (based on GBP1 (PDB: 1f5n) ^41^.

All further analysis was performed in UCSF Chimera ^42^. The structures of GBP1 and the predicted structures were aligned using the “MatchMaker” function and critical amino acids of the DFCP1 P-Loop domain were identified by their structural proximity to the corresponding amino acids of GBP1. The structural alignment was also used to approximate the localization of the bound nucleotide, shown is the GDP moiety bound by GBP1.

For further comparison to other nucleotide-binding proteins, we generated a structural alignment using the “Match-Align” feature of Chimera, using DFCP1, hGBP1 (PDB:1f5n), hAtlastin1 (PDB: 3q5e), N-RAS Q61H (PDB: 621p) and KRas (PDB:4obe) as templates.

### Protein purification

Recombinant proteins were expressed as His-tagged MBP fusion proteins in E. coli. Constructs encoding the DFCP1 N-terminus were cloned into pET-His6-MBP-TEV-LIC (Addgene # 29708) and Rosetta2(DE3) bacteria were transformed with the resulting plasmids. For protein expression, bacteria were grown in ZYM505 medium ^43^. Expression was induced by 0.25 mM IPTG, and induced cells were grown over night at 20°C. Cells were harvested by centrifugation, resuspended in lysis buffer (50 mM Tris pH7.5, 150 mM NaCl, 10 mM MgCl_2,_ 25 mM Imidazole, 1 mM TCEP, cOmplete mini EDTA-free protease inhibitor (Roche)) and lysed by one passage through a homogenizer. Raw lysates were cleared by centrifugation. The His-MBP fusion proteins were purified by affinity chromatography using NiNTA affinity chromatography. Protein-containing fractions were pooled, dialyzed over night against dialysis buffer (50 mM Tris pH7.5, 150 mM NaCl, 10 mM MgCl_2,_ 10% Glycerol, 1 mM TCEP). Aliquots were snap-frozen in liquid nitrogen and stored at −80°C.

### Nucleotide binding assays

Nucleotide binding assays were carried out by measuring the incorporation of 2,3-0-N-Methyl-anthraniloyl (Mant)-labelled nucleotides (Jena Bioscience) using an Synergy2 plate reader (BioTek Instruments Inc., Winooski, VT, USA) at 25°C. Reactions containing 1 µM mantNTP, 50 mM Tris pH7.5, 150 mM NaCl, 50 mM MgCl_2,_ 10% Glycerol, 100 µg/ml BSA were allowed to equilibrate for 4 mins. After equilibration, recombinant proteins were added to a final concentration of 2µM and the fluorescence intensity (λ_ex_=360 nm, λ_em_=440nm) was measured.

### Measurement of ATPase activity

ATPase activity was measured by quantifying the release of inorganic phosphate using a modified Malachite Green Phosphate assay (Biomol Green, Enzo). All measurements were performed according to the manufacturer’s protocol. Briefly, 10µM purified DFCP1 N-terminus was incubated with 500 µM ATP for 30min. After 30 min, two volumes of Biomol Green were added and incubated for further 20 min before the OD620 was recorded.

### Size exclusion chromatography

Size exclusion chromatography was performed using an Äkta Explorer (Cytiva) on a Superdex 200 Increase 10/300 GL column (Cytiva). Recombinant DFCP1 was loaded with nucleotides by incubation with either 400 µM ADP or ATPγS (Jena Bioscience) in exchange buffer (50 mM Tris, 150 mM NaCl, 50 mM MgCl_2_, 10% Glycerol, 1 mM TCEP) for 20 min at RT, applied to the size exclusion column and then developed by isocratic elution with exchange buffer at 4°C (flow rate 0.25 ml/min).

### Cell culture

U2OS cells (ATCC: HTB-96), HeLa cells (obtained from Institute Curie, Paris, France), and A431 cells were maintained in DMEM (D0819; Sigma-Aldrich) supplemented with 10% FCS (F7524; Sigma-Aldrich), 5 U/ml penicillin and 50 μg/ml streptomycin at 37°C with 5% CO_2_. hTERT-RPE-1 cells (ATCC: CRL-4000) were grown in DMEM/F12 medium (31331–028; Gibco-BRL) with 10% FCS, 5 U/ml penicillin and 50 µg/ml streptomycin at 37°C with 5% CO_2_. Parental cell lines and its derivatives were regularly tested for mycoplasma. For starvation experiments, cells were washed twice with EBSS (24010-043; GIBCO BRL) and incubated in EBSS for the indicated time. Bafilomycin A1 (Enzo life sciences, BML-CM-110-0100) was added at 100 nM, Puromycin at 2.5-10 µg/ml (Sigma, P8833), DFP (#379409, Sigma-Aldrich) at 0.5-1 mM, and Torin1 (#4247, Bio-Techne R&D systems Europe) at 50 nM.

### Generation of cell lines

Cell lines stably expressing constructs were generated lentiviral transduction using a third-generation lentiviral vector system as previously described ^44^. Subsequently, cells were selected for integration of the expression cassette with the following antibiotics: Puromycin (1 μg/ml), Blasticidin (10 μg/ml), or Geneticin (500 μg/ml). All lentiviral constructs were expressed from a phospho-glycerate kinase (PGK) promoter.

### Generation of DFCP1 KO cell line with CRISPR/Cas9

Deletion of DFCP1 in parental U2OS and A431 cells was generated using the CRISPR/Cas9 system. Two guide RNAs were used due to the presence of internal start codons in Exons 5-7 after the primary start codon in Exon2. The guide RNAs were designed in Benchling (http://www.benchling.com). gRNA1 ‘5-TGTCCCTTACTGTGACCTCT-3’ binds after the primary start codon in Exon2, gRNA2 ‘5-TGGCGTGGTCTATCGTAGT-3’ binds in Exon8. The guide RNAs were cloned into a pX459 plasmid encoding Cas9-2A-Puro, and the final construct was transfected into the cells with FugeneHD (U20S) or Lipofectamine LTX with Plus reagent (A431). After 48 h post-transfection cells were selected with Puromycin. Single colonies were picked and characterized. Clones lacking DFCP1 were identified by western blotting, and the genetic changes were characterized by PCR and sanger sequencing. Genetic changes sequences were analysed by using the Synthego ICE software package (https://ice.synthego.com/#/).

### Generation of DFCP1 KI cell line with CRISPR/Cas9

U2OS cells expressing endogenously tagged DFCP1 were generated by using CRISPR/Cas9 in conjunction with an AAV-based homology donor. Cells were first transduced with the AAV harbouring the homology donor, and electroporated 16 h later with CRISPR/Cas9 RNP particles using a Nepa21 electroporator. Recombinant Cas9, crRNA and tracrRNA were purchased from IDT and RNP complexes were formed according to the manufacturer’s manual. The gRNA was located at the start-codon of DFCP1 (gRNA sequence: CTGGGCACTCATACTCACGC).

The tagging strategy was based on a two-step tagging technique ^45^ using a tagging cassette based on the plasmid FLAG-LoxP-PURO-LoxP-HaloTERT WT HR Donor (Addgene # 86843), with the Halo-Tag exchanged by EGFP. This technique first integrates a tagging construct including a resistance marker, which is excised by Cre-mediated recombination in a second step. To generate the homology donor, a construct containing ∼1KB of homology left and right of the start codon and the tagging was assembled in an AAV vector (pAAV-2Aneo v2, Addgene plasmid 800333) and packaged using pHelper and pRC2-miR321 vectors (Clontech). AAV particles were isolated using AAVPro extraction solution (Clontech). After transduction with both CRISPR/Cas9 gesicles and AAV homology donor, cells with integrated homology donor were selected using puromycin selection (1 µg/ml). Single clones were picked and characterized by PCR and western blotting and the integration site verified by sequencing. Successful homozygous clones were then treated with Cre recombinase to excise the resistance marker, resulting in a cell line expressing FLAG-EGFP-DFCP1 under control of the endogenous promoter.

### siRNA-mediated knockdown

For siRNA transfections, 20 nM of siRNA was transfected using Lipofectamine RNAiMax reagent. Transfected cells were analysed 48 h to 72 h after transfection; knockdown efficiency was verified by western blotting. For knockdown of DFCP1 or ATG13, the following Silencer Select siRNAs from Ambion (Thermo Fisher Scientific, Waltham, MA, USA) were used: Silencer^®^ Select Negative Control No.1 (4390843), siDFCP1-1 (ID s28712, siDFCP1-2 (ID s28714), siDFCP1-3 (ID 135291), or siATG13 (ID s18879).

### Immunoblotting

Cells were washed with ice-cold PBS and lysed in 2.8x Laemmli Sample buffer (BioRad) containing 200 mM DTT. Cell lysates were subjected to SDS-PAGE on a 4–20 % (567–1094; Bio-Rad) gradient gel and blotted onto PVDF membranes (BioRad). Membranes incubated with fluorescently labelled secondary antibodies (IRDye680 and IRDye800; LI-COR) were developed by Odyssey infrared scanner (LI-COR). Membranes detected with HRP labelled secondary antibodies were developed using Clarity Western ECL substrate solutions (Bio-Rad) with a ChemiDoc XRS+ imaging system (Bio-Rad).

### Co-immunoprecipitation

HeLa cells were transfected with pEGFP-N1 or pEGFP-DFCP1 and co-transfected with pcDNA3-myc-Ubiquitin using Fugene6 (Promega 2023-11-06 Lot 0000425615). 24 h later the cells were washed twice in ice cold PBS and lysed (125 mM KAc, 25 mM Hepes, 2.5 mM MgAc (4*H_2_O), 5 mM EGTA, pH 7.2, substituted freshly with complete protease inhibitor (Roche), PhosSTOP phosphatase inhibitor (Roche), 0.5 % IGEPAL and 1 mM DTT). Lysates were centrifuged for 5 min at 20 000 x g and supernatants were immunoprecipitated for 1 h rotating at 4°C with mouse anti-GFP antibody (1 µg antibody per sample, 11814460001 Merck/Sigma) and 20 µl DynabeadsTM Protein G (10004D, Thermo Fisher). The immunoprecipitates were washed three times in lysis buffer and eluted with 2x Laemmli sample buffer (BioRad) containing DTT and analysed by immunoblotting.

### Measuring autophagy by ratiometric flow cytometry

The monomeric Keima (mKeima) protein is a useful tag for measuring autophagic degradation of various autophagic cargoes, such as mitochondria ^19^. Since mKeima is resistant to lysosomal proteases and has a bimodal, pH-responsive excitation spectrum, the assay provides a cumulative readout of autophagic activity as mKeima stably accumulates and undergoes a change in excitation maximum upon trafficking to the acidic environment of lysosomes ^20, 46^. A probe for mitophagy was obtained by fusing the mitochondrial-targeting pre-sequence of COX VIII to mKeima, producing mito-mKeima. To compare mitophagy with bulk autophagy of cytosolic proteins, we used either free mKeima itself or the cytosolic protein lactate dehydrogenase (LDHB) fused to mKeima as cytosolic probes ^19^. RPE-1 cells expressing LDHB-mKeima or mito-mKeima ^26^ and U2OS cells expressing free mKeima or mito-mKeima in an inducible manner were generated by lentiviral transduction. The cells were treated with doxycycline (200 ng/ml) to induce expression of the mKeima probes during the 2 days of siRNA-mediated depletion of DFCP1 or ATG13. Before further treatments, the cells were washed twice to remove doxycycline. The cells were subsequently treated with EBSS, Torin1 (50 nM) or DFP (0.5 mM) for 16 h, detached by trypsin/EDTA and resuspended in complete cell culture medium for analysis on a BD LSR II Flow Cytometer (BD Biosciences, San Jose, CA, USA) connected to the BD FACSDiva™ software (BD Biosciences, San Jose, CA, USA). Autophagy was determined as the ratio of the median fluorescent intensity of the mKeima signal excited by 561 nm (45 mW) laser divided by mKeima excited by 407 nm (100 mW) laser, with a 610/20 bandpass filter and a 600 nm long pass dichroic filter in both cases. By using the FlowJo™ software (BD Biosciences, San Jose, CA, USA), the derived ratio of 561/407 nm signal intensity per cell was obtained, and the median value of these cellular ratios were compared between treatments.

### Puromycin treatment to induce aggrephagy

Puromycin (Sigma, P8833) was added at a concentration of 2.5 or 10 µg/ml to complete medium, cells were incubated for the indicated time, fixed and stained as described below.

### Immunofluorescence microscopy and image analysis

Cells were seeded on glass coverslips, fixed with 4% formaldehyde (FA; 18814; Polysciences) for 12 min at room temperature, and permeabilized with 0.05% saponin (S7900; Sigma-Aldrich) in PBS. Fixed cells were then stained with primary antibodies at room temperature for 1 h in PBS/saponin, washed in PBS/saponin, stained with fluorescently labelled secondary antibody for 1 h, washed in PBS, and mounted with Mowiol containing 2 µg/ml Hoechst 33342 (H3570; Thermo Fisher Scientific).

Confocal micrographs were obtained using an LSM710 confocal microscope (Carl Zeiss) equipped with an Ar-laser multiline (458/488/514 nm), a DPSS-561 10 (561 nm), a continuous-wave laser diode 405–30 CW (405 nm), and an HeNe laser (633 nm). The objective used was a Plan-Apochromat 63×/1.40 oil differential interference contrast (DIC) III (Carl Zeiss). For visualization, images were analysed and adjusted (brightness/contrast) in ImageJ/Fiji ^47^ or Zen Blue.

For quantification, all images within one dataset were taken at fixed intensities below saturation, very high expressing cells have been omitted, and identical settings were applied for all treatments within one experiment. In general, 25 images were taken randomly from each condition.

The NIS-elements software was used for background correction (“rolling ball”) and automated image analyses. Identical analysis settings were applied for all treatments within one experiment. Fluorescent LC3B, p62 or WIPI2 dots were segmented by the software, and the number and sum fluorescence intensity of the dots were measured, as well as the co-occurrence of WIPI2 and DFCP1 dots. The total number of cells was quantified by automated detection of Hoechst nuclear stain by the software. The number of mNG-DFCP1 omegasome rings was quantified manually, scoring all objects with a visible lumen. Micronuclei were quantified manually from micrographs.

### Live Imaging of omegasome formation

Live-cell imaging was performed on a Deltavision OMX V4 microscope equipped with three PCO.edgesCMOS cameras, a solid-state light source and a laser-based autofocus. Environmental control was provided by a heated stage and an objective heater (20–20 Technologies). Images were deconvolved using softWoRx software and processed in ImageJ/FIJI. Cells were imaged in EBSS, and images were taken every 2 sec over a period of 10-15 min. For imaging of SNAP-LC3B, cells were stained with SiR647-SNAP (New England Biolabs) according to the manufacturer’s protocol.

### Structured illumination microscopy (SIM)

Cells were prepared according to the immunofluorescence protocol detailed above. Three-dimensional SIM imaging was performed on Deltavision OMX V4 microscope with an Olympus ×60 NA 1.42 objective and three PCO.edge sCMOS cameras and 488 nm, 568 nm and 647 nm laser lines. Cells were illuminated with a grid pattern and for each image plane, 15 raw images (5 phases and 3 orientations) were acquired. Super-resolution images were reconstructed from the raw image files and aligned using softWoRx software (Applied Precision, GE Healthcare). Images were processed in ImageJ/Fiji. 3D reconstructions were performed using Imaris.

### Statistics

The number of individual experiments and the number of cells or images analysed are indicated in the figure legends. The number of experiments was adapted to the expected effect size and the anticipated consistency between experiments. We tested our datasets for normal distribution by Kolmogorov-Smirnov, D’Agostino and Pearson, and Shapiro–Wilk normality tests, using GraphPad Prism Version 8. For parametric data, an unpaired two-sided t test was used to test two samples with equal variance, and a one-sample t test was used in the cases where the value of the control sample was set to 1. For more than two samples, we used ordinary one-way ANOVA with a suitable post hoc test. For nonparametric samples, Mann–Whitney test was used to test two samples and Kruskal–Wallis with Dunn’s post hoc test for more than two samples. All error bars denote mean values ±SD or SEM, as indicated in every figure legend (*, P < 0.05; **, P <0.01; ***, P < 0.001). No samples were excluded from the analysis.

### Movie legends

#### Movie 1, corresponding to Fig.2b

U2OS DFCP1 KO lines stably expressing mNG-DFCP1 WT and mCh-p62. Cells were starved in EBSS and imaged live to capture omegasome formation. Omegasome formation was divided into three phases as indicated in Fig2a. Quantification in Fig.2b, c. Frame 2 sec. Scale bar 1 µm.

#### Movie 2, corresponding to Extended Data Fig.4a

U2OS DFCP1 KO lines stably expressing mNG-DFCP1 WT or -mutant and mCh-p62. Cells were starved with EBSS and imaged live to capture omegasome formation. Quantification in Fig.2b,c. Frame 2 sec. Scale bar 1 µm.

#### Movie 3, corresponding to Extended Data Fig.4d

U2OS DFCP1 knockdown/rescue lines stably expressing mNG-DFCP1 WT and SNAP-LC3B were depleted by endogenous DFCP1 for 2 d, starved with EBSS and imaged live to capture omegasome formation. Omegasome formation was divided into three phases as indicated in Extended Data Fig.3d. Frame 2 sec. Scale bar 1 µm.

#### Movie 4, corresponding to Extended Data Fig.4f

U2OS DFCP1 knockdown/rescue lines stably expressing mNG-DFCP1 WT or -mutant and SNAP-LC3B were depleted by endogenous DFCP1 for 2 d, starved with EBSS and imaged live to capture omegasome formation. Frame 2 sec. Scale bar 1 µm.

#### Movie 5, corresponding to Fig.4d

Example of a mitophagy event. U2OS cells stably expressing mNG-DFCP1 WT and SNAP-mito were treated for 18 h with 0.5 mM DFP and imaged live. The movie shows a piecemeal of mitochondria, which is engulfed by the phagophore. Note that the phagophore is weakly positive for mNG-DFCP1. Frame 2 sec. Scale bar 1 µm.

#### Movie 6, corresponding to Extended Data Fig.8d

Example of a mitophagy event. U2OS cells stably expressing mNG-DFCP1 WT and SNAP-mito were treated for 18 h with 0.5 mM DFP and imaged live. Note the lumen of the omegasome ring, through which a part of the mitochondria is threaded. Frame 2 sec. Scale bar 1 µm.

## Acknowledgements

We thank the Flow Cytometry Core Facility and the Advanced Light Microscopy Core Facility of Oslo University Hospital, and the Helsinki Institute of Life Science (HiLIFE) Light Microscopy Unit for technical assistance and access to instruments. We thank Ulrikke Dahl Brinch for general technical help and the entire Stenmark group for discussion.

## Author Contributions

V.N and K.O.S. conceived and designed the study with contributions from C.R. and H.S. V.N. generated cell lines, performed light microscopy and biochemical assays, analysed data, prepared figures, and wrote the original draft. K.O.S. performed structural analyses, biochemical assays, prepared figures and edited the draft. C.R. performed HiC imaging, analysed data, prepared figures, and edited the draft. K.W.T. generated cell lines, performed HiC imaging, analysed data, and prepared figures. M.N. and T.J. performed and analysed GST binding assays, prepared figures and edited the draft. M.L.T. performed the mKeima assays, analyzed data and prepared figures. S.M. generated KO and KI lines, performed SIM, analysed data, and prepared figures. E.M.W. performed immunoprecipitation experiments, analysed data, and prepared figures. V.S. and E.I. performed and analysed experiments in A431 cells and edited the manuscript. H.S. supervised the study, contributed to funding acquisition and research infrastructure, and edited the manuscript.

## Funding

V.N. was supported by a Mobility grant from the Research Council of Norway (grant number 301369). C.R. was supported by a Research Grant from the Norwegian Cancer Society (grant number 198140). M.L.T. was supported by the Research Council of Norway (grant number 274574). V.T.S. was supported by Biomedicum Helsinki Foundation. M.N. was supported by a Grant from the Research Council of Norway (grant number 249884) to T.J. E.M.W was supported by the Radium Hospital Foundation (InvaCell grant / Trond Paulsen). K.O.S was supported by a Career grant from the South-Eastern Norway Regional Health Authority (grant number 2020038) and a Research Grant from the Research Council of Norway (grant number 315103). H.S. was supported by Project Grants from the South-Eastern Norway Regional Health Authority (project number 2018081) and the Norwegian Cancer Society (project number 182698), and by an Advanced Grant from the Europan Research Council (project number 788954). This work was partly supported by the Research Council of Norway through its Centres of Excellence funding scheme, project number 262652.

## Competing interests

The authors declare no competing financial interests.

---

**Extended Data Fig.1:**
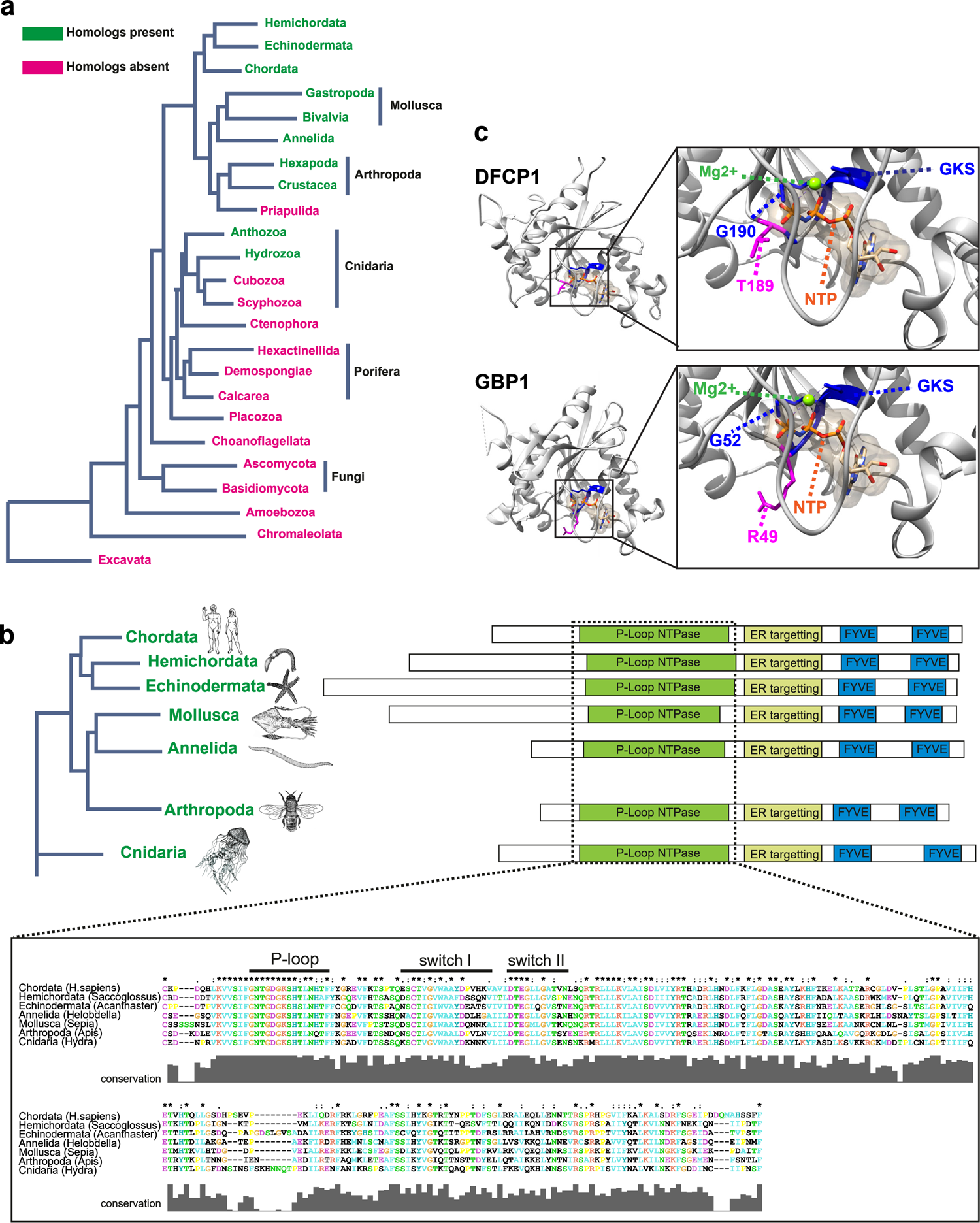
DFCP1 is an ancient protein. (a) DFCP1 is conserved in the animal kingdom. Homologs of DFCP1 first occur in the cnidarian phylum and is present in nearly all metazoan phyla, indicating that it was first invented more than 0.5 billion years ago. (b) The domain structure of DFCP1 is highly conserved throughout the metazoan tree of life. The DFCP1 domain architecture has remained largely unchanged from the first occurrence in the cnidarians. All homologs have a highly conserved ATP binding domain, an ER-binding domain and two FYVE domains. (c) Phyre2-generated homology model of DFCP1, based on the structure of GBP1. Candidates for amino acids required for nucleotide binding (GKS, residues 192-194) are indicated in blue, and amino acids at the same position as the catalytic arginine of GBP1 (T189, G190) and KRAS (G12, G13) are indicated in magenta.

**Extended Data Fig.2:**
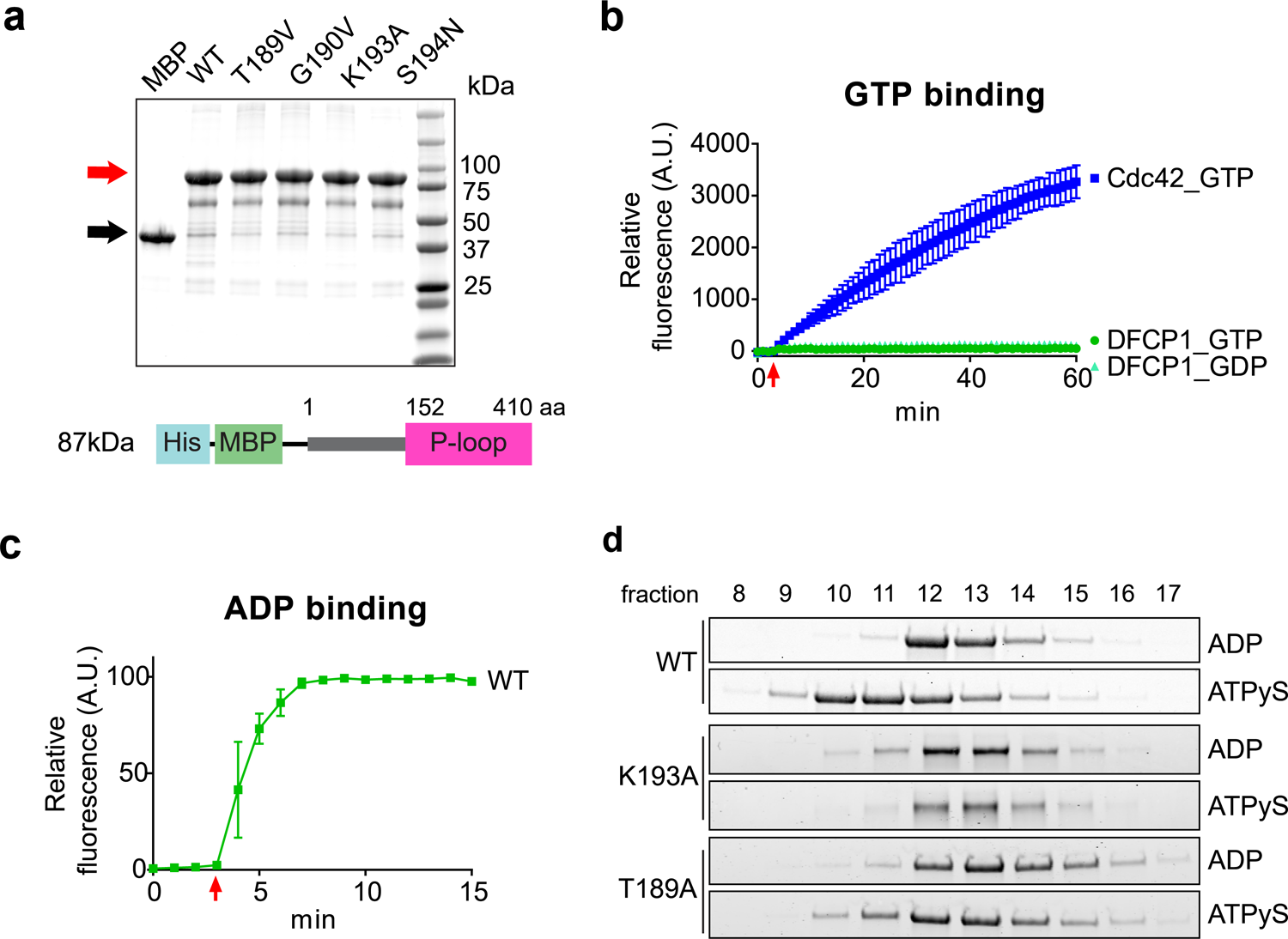
DFCP1 binds and hydrolyses ATP. (a) Coomassie gel from purified DFCP1 N-terminus as used in Fig.1d-h and Extended Data Fig.2b-d. Red arrow indicates MBP-DFCP1 for a calculated mass of 87kDa, black arrow indicates MBP (43 kDa). Schematic overview of the His- and MBP-tagged N-terminus including P-loop of DFCP1 (DFCP1 K193A or DFCP1 T189A) used for nucleotide binding measurements. (b) DFCP1 does not bind guanosine nucleotides. Recombinant DFCP1 WT was incubated in the presence of mantGTP (n=4) or mantGDP (n=3) (arrow). Nucleotide binding was measured by incorporation of fluorescent mantGDP/GTP. Purified Cdc42 was used as a positive control for GTP binding (n=2). Curves are normalized to MBP (DFCP1) or GST (Cdc42) background fluorescence. Bars: mean ± SD. (c) DFCP1 binds to adenosine nucleotides. DFCP1 WT was incubated with mantADP (arrow), and nucleotide binding was measured by the incorporation of fluorescent mantATP/ADP. Curve is normalized to MBP background fluorescence. Bars: mean ± SD, 3 experiments. (d) Coomassie gel showing individual 1 ml fractions of the size exclusion chromatography shown in Fig.1h.

**Extended Data Fig.3:**
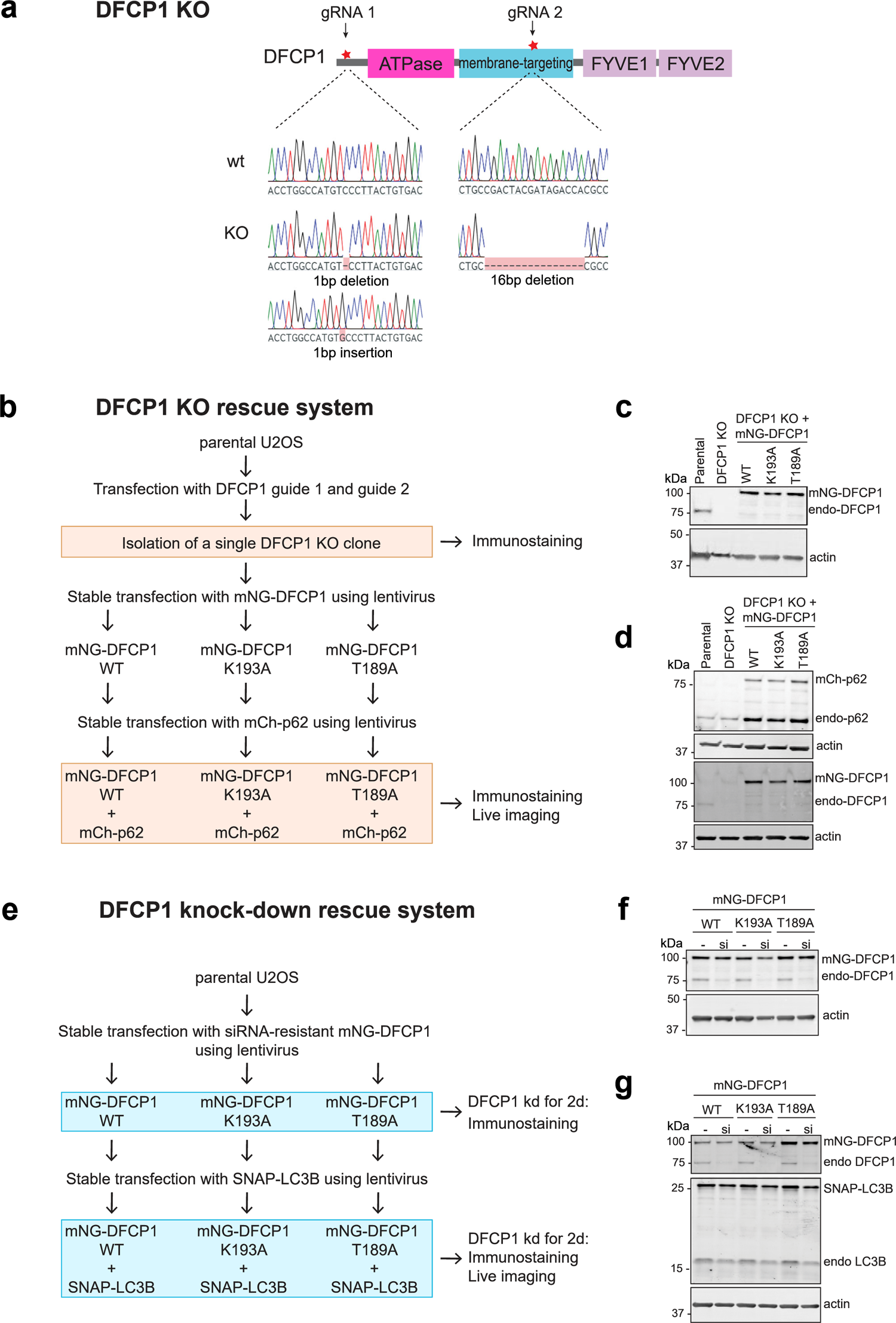
Characterization of DFCP1 KO- and knockdown-rescue systems. (a) Generation and characterization of CRISPR/Cas9-mediated DFCP1 KO in U2OS cells. Shown are the localization of guide RNAs targeting DFCP1 and the sequences of a single clone which was isolated and validated by Sanger sequencing. This clone has been utilized for all further experiments. (b) Schematic drawing of the generation of the DFCP1 KO rescue system. DFCP1 KO cells were stably transduced with a low titer of mNG-DFCP1 WT, K193A or T189A. These lines were further transduced with mCh-p62, and cells have been imaged live in EBSS for omegasome quantification (Fig.2, Extended Data Fig.4a-c). (c) Western blotting was used to measure protein levels of endogenous DFCP1 and mNG-DFCP1 using anti-DFCP1 antibody. Actin was used as loading control. (d) Western blotting was used to measure protein levels of endogenous and exogenous expressed DFCP1 and p62 using anti-DFCP1 and anti-p62 antibodies. Actin was used as loading control. (e) Schematic drawing of the generation of the DFCP1 knockdown-rescue system for confocal imaging. U2OS cells were stably transduced with a low titer of siRNA-resistant mNG-DFCP1 WT, K193A or T189A. The resulting stable cell lines were then transfected with siRNA against DFCP1 for 2 d and treated as mentioned. Alternatively, cells were transduced with lentiviral vectors expressing SNAP-LC3B and cells were imaged live in EBSS for omegasome quantification (Extended Data Fig4.d-i). (f) Western blotting was used to validate knockdown of DFCP1 using anti-DFCP1 antibody. Actin was used as loading control. (g) Western blotting was used to measure protein levels of DFCP1 and LC3B, actin was used as loading control.

**Extended Data Fig.4:**
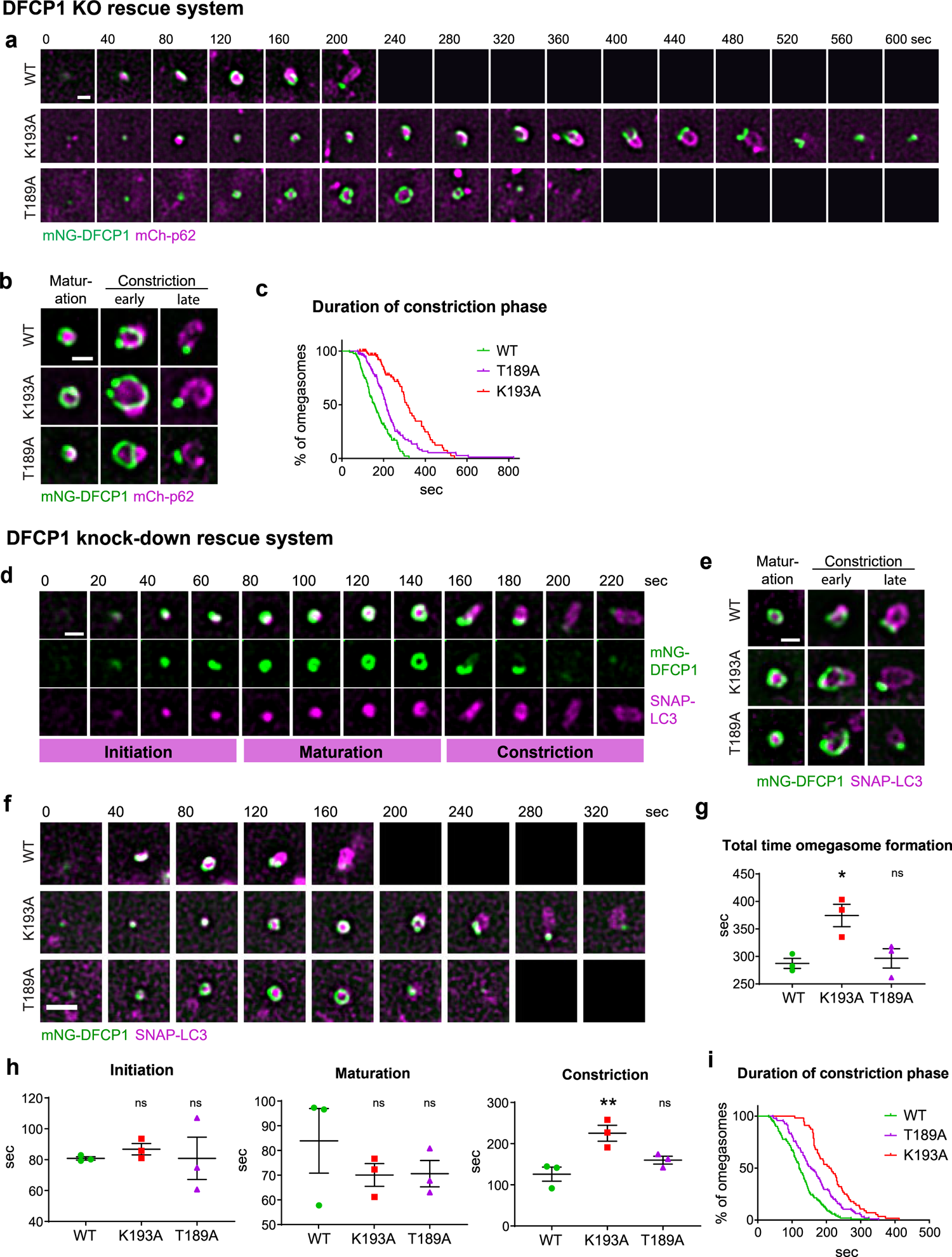
DFCP1 mutants are delayed in omegasome constriction. (a) U2OS DFCP1 KO lines stably expressing mNG-DFCP1 WT or -mutant and mCh-p62. Cells were starved with EBSS and imaged live to capture omegasome formation. Shown are representative stills of omegasomes analyzed in Fig.2b, c. Scale bar 0.5 µm. (b) Examples of omegasomes from cells expressing DFCP1 WT or mutant DFCP1. U2OS DFCP1 KO lines stably expressing mNG-DFCP1 WT and mCh-p62 were starved in EBSS and imaged live to capture omegasome formation. The first image shows the ring phase, with p62 localizing in the middle. The second image shows the time point which was used as the onset of the constriction phase. A p62-positive autophagosome with a remaining DFCP1 dot is shown in the third image. Scale bar 0.5 µm. Representative image of > 20 events. (c) Duration of the constriction phase of single omegasomes plotted over time. The quantification is derived from the same raw data as in Fig.2b-d. (d) U2OS DFCP1 knockdown/rescue lines stably expressing mNG-DFCP1 WT and SNAP-LC3B were depleted by endogenous DFCP1 for 2 d, starved with EBSS and imaged live to capture omegasome formation. Omegasome formation was divided into three phases as indicated. Shown are representative stills of omegasomes analyzed in (g) and (h). Scale bar 0.5 µm. (e) Examples of omegasomes from DFCP1 knockdown rescue lines expressing mNG-DFCP1 WT or - mutant and SNAP-LC3B. The first image shows the ring phase, with SNAP-LC3B localizing in the middle. The second image shows the time point which was defined as the onset of the constriction phase. A LC3B-positive autophagosome with a remaining DFCP1 dot is shown in the third image. Scale bar 0.5 µm. Representative image of > 40 events. (f) U2OS DFCP1 knock-down rescue lines stably expressing mNG-DFCP1 WT or -mutant and SNAP-LC3B were depleted by endogenous DFCP1 for 2 d, starved with EBSS and imaged live to capture omegasome formation. Scale bar 0.5 µm. (g) Total time of an omegasome life cycle, with a DFCP1 punctum as beginning and -end points. Each plotted point represents the mean value of one experiment. Bars: mean ± SEM, 3 experiments. Statistics: One-way ANOVA followed by Dunnett’s multiple comparisons test, comparing against WT. Analyzed omegasomes per genotype: WT=93, K193A=45, T189A=89. (h) Duration of the individual phases of omegasome formation. Each plotted point represents the mean value of one experiment. The quantification is derived from the same raw data as (f) and (g). Bars: mean ± SEM, 3 experiments. Statistics: One-way Anova followed by Dunnett’s multiple comparisons test, comparing against WT. Analyzed omegasomes per genotype and phase: Initiation: WT=77, K193A= 48, T189A=76; Maturation: WT=87, K193A=56, T189A=83; Constriction: WT=103, K193A=56, T189A=92. (i) Duration of the constriction phase of single omegasomes plotted over time. The quantification is derived from the same raw data as in (g) and (h).

**Extended Data Fig.5:**
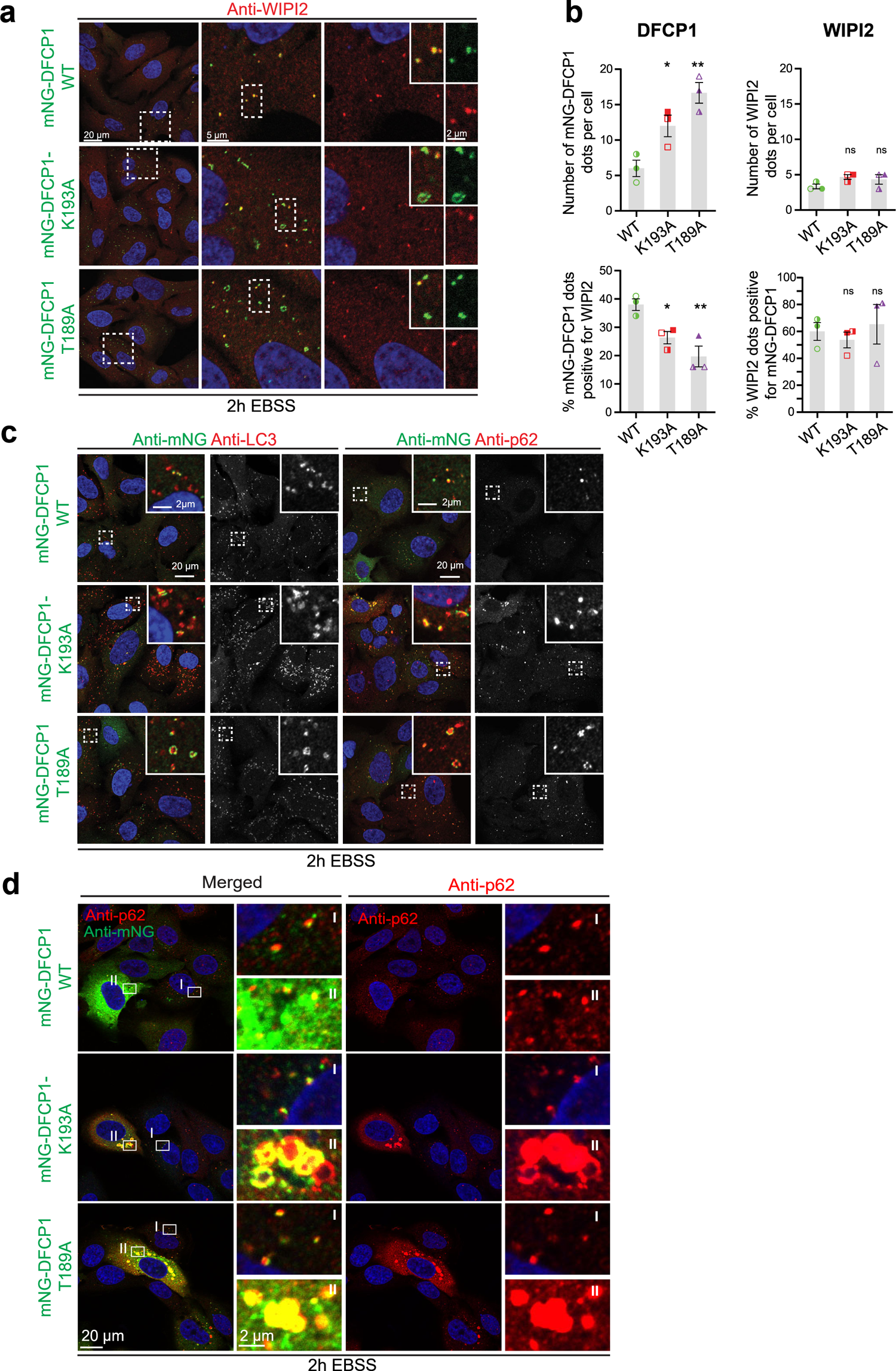
Omegasome initiation is not affected in DFCP1 ATPase mutants. Fixed imaging of U2OS DFCP1 knock-down rescue lines stably expressing mNG-DFCP1 WT, K193A or T189A as indicated: (a) Confocal images showing the localization and abundance of WIPI2 puncta in DFCP1 mutant cells. U2OS cells stably expressing siRNA resistant mNG-DFCP1 WT, K193A or T189A, were depleted of endogenous DFCP1 by siRNA transfection for 2 d in complete medium before starvation for 2 h in EBSS. The cells were fixed, stained with anti-WIPI2 antibody, and analyzed by confocal microscopy. Insets show that some of the DFCP1-positive omegasomes co-localize with WIPI2. Representative image for (b). Scale bar 20 µm, insets 5 µm and 2 µm. (b) Graphs represent quantifications of (a) of the amount of mNG-DFCP1 dots and WIPI2 dots per cell and the co-occurrence between DFCP1 and WIPI2 dots, from images like described above. Error bars denote mean ± SEM, 3 experiments. Each plotted point represents the mean value of one experiment, and independent experiments are indicated by different colored or transparent objects. Ordinary one-way ANOVA, Dunnett’s post hoc test: * p < 0.05; ** p < 0.01; ns: not significant. Analyzed cells per condition: WT= 268, K193A= 273, T189A= 227. (c) Representative images of the quantifications shown in Fig.3c. Confocal images showing the localization of DFCP1, DFCP1 mutants, LC3B and p62 in DFCP1 KD rescue cell lines. U2OS cells stably expressing siRNA resistant mNG-DFCP1-WT, K193A, or T189A were depleted of endogenous DFCP1 by siRNA transfection for 2 d in complete medium before starvation for 2 h in EBSS. Cells were fixed, stained with anti-mNG and either anti-LC3B or anti-p62 antibodies and analyzed by confocal microscopy. Both LC3B and p62 are found in close association with DFCP1 positive omegasomes. Note that the localization of p62 is more restricted to omegasomes than LC3B. Quantifications in Fig3c, d. Scale bar 20 µm, inset 2 µm. (d) U2OS cells stably expressing DFCP1 WT, K193A, T189A were depleted of endogenous DFCP1 by siRNA, treated with EBSS for 2 h, fixed and stained with anti-mNG and anti-p62 antibodies and analyzed by confocal microscopy. p62 colocalizes with DFCP1 in high and low expressing DFCP1 cells under all conditions. Notably, strong overexpression of the ATPase defective DFCP1 T189A allele causes hyper-accumulation of p62. Such cells, which appeared with low frequency in the stable cell lines, were not included in the high content-based image quantifications. Representative of >5 images per condition.

**Extended Data Fig.6:**
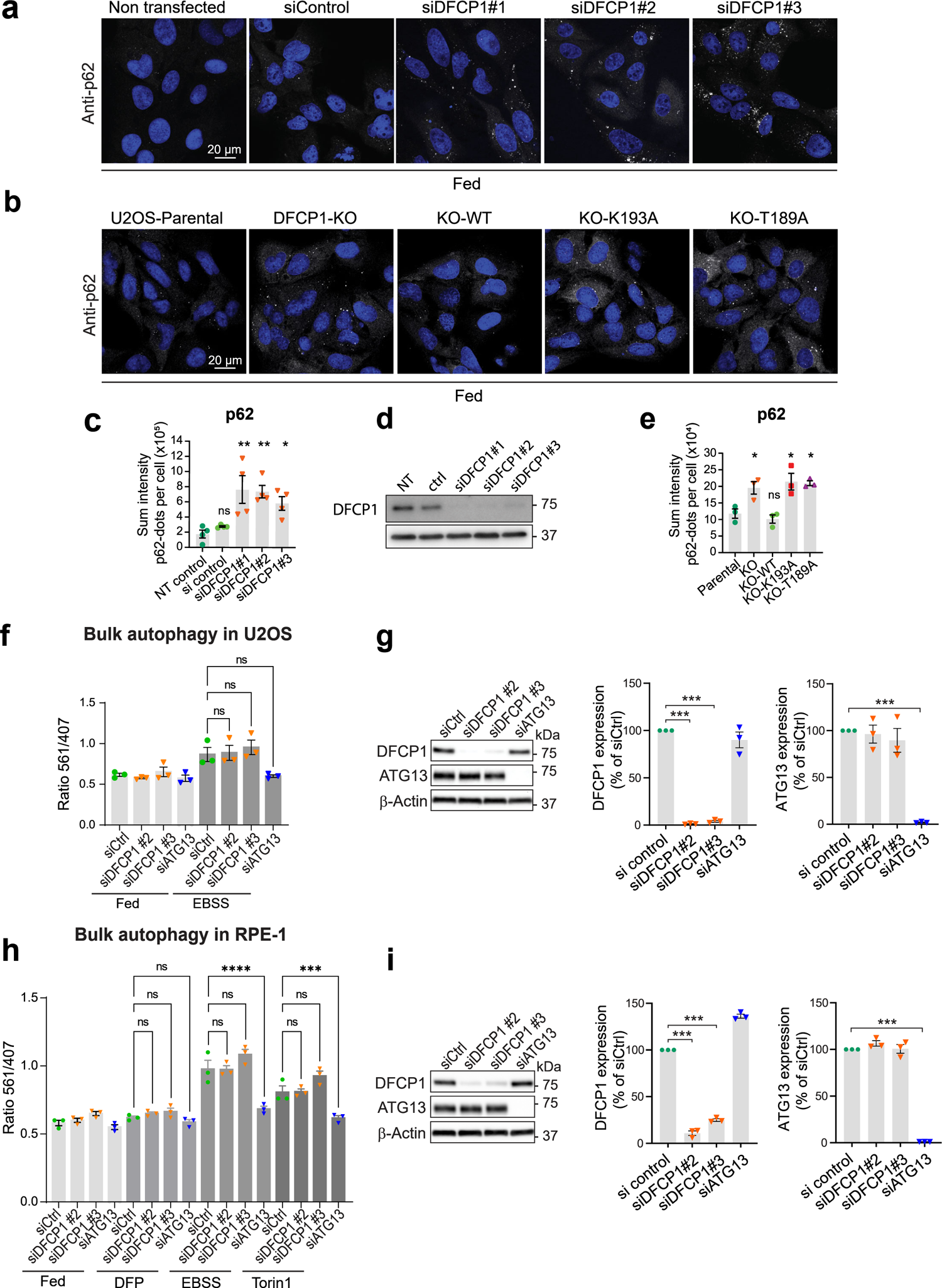
While bulk autophagy is not affected, the levels of p62 are elevated in DFCP1 ATPase mutants. (a) U2OS cells were depleted of endogenous DFCP1 by siRNA transfection for 2 d in complete medium, fixed, stained with anti-p62 antibody and Hoechst33342, and analyzed by confocal microscopy. Scale bar 20 µm. (b) U2OS parental cells and DFCP1 KO rescue cells cultured in complete medium, were fixed and stained with anti-p62 antibody and Hoechst33342 and analyzed by confocal microscopy. Scale bar 20 µm. (c) Quantifications of (a). Graphs represent quantification of the sum intensity of p62 dots per cell. Error bars denote mean ±SEM, 4 experiments. Each plotted point represents the mean value of one experiment. Ordinary one-way ANOVA, Dunnett’s post hoc test: * p < 0.05; ** p < 0.01; ns: not significant. Total cells analyzed per condition: NT (non-transfected): 575, siCtr= 507, si#1= 464, si#2= 504, si#3= 507. (d) Western blotting was used to validate knockdown of DFCP1 in (a) using anti-DFCP1 antibody. Actin was used as loading control. Representative blot from n=4 experiments. (e) Quantification of the experiment shown in (b). Graphs represent quantification of the sum intensity of p62 dots per cell. Error bars denote mean ± SEM from n=3 independent experiments. Each plotted point represents the mean value of one experiment. Ordinary one-way ANOVA, Dunnett’s post hoc test: * p < 0.05; ns: not significant. Cells analyzed per condition: parental: 788; KO:703; WT: 732, K193A: 779, T189A: 766. (f) U2OS cells induced to express free cytoplasmic mKeima were transfected with control-, DFCP1- or ATG13 siRNA for 2 d. Cells were either grown for 16 h in complete medium or starved in EBSS. Cells were harvested and subjected to Flow Cytometry to determine the ratio of lysosome-localized mKeima to free, cytoplasmic mKeima. Bars: ±SEM, 3 experiments. Statistics: One-way Anova followed by Dunnett’s multiple comparisons test. (g) Knockdown validation for (f) with Western Blot and quantification of DFCP1- and ATG13 bands using Actin as a loading control. Values were normalized to control siRNA. Bars: ±SEM, 3 experiments. Statistics: Welch’s test, comparing against siCtrl. (h) RPE-1 cells induced to express LDHB-mKeima were transfected with control-, DFCP1- or ATG13 siRNA for 2 d. Cells were cultured for 16 h either in complete medium, in 0.5 mM DFP, were starved in EBSS, or treated with 50 nM Torin1. Cells were harvested and subjected to Flow Cytometry to determine the ratio of lysosome-localized LDHB-mKeima to free, cytoplasmic LDHB-mKeima. Bars: ± SEM, 3 experiments. Statistics: One-way Anova followed by Dunnett’s multiple comparisons test. (i) Knockdown validation for (h) with Western Blot and quantification of DFCP1- and ATG13 bands using Actin as a loading control. Values were normalized to control siRNA. Bars: ±SEM, 3 experiments. Statistics: One sample t-test, comparing against a theoretical mean of 100.

**Extended Data Fig.7.**
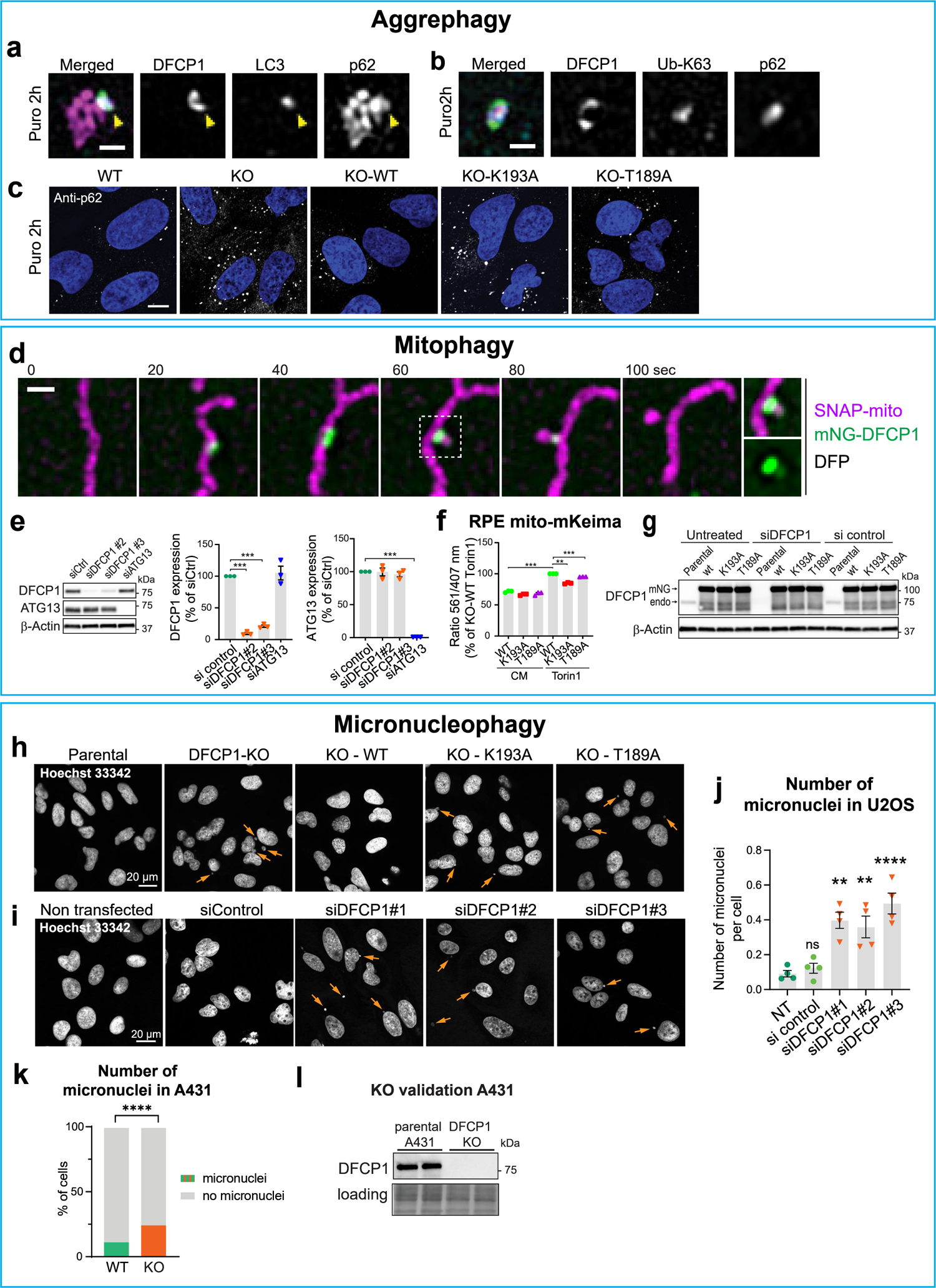
DFCP1 ATPase regulates selective types of autophagy. (a) U2OS stably expressing mNG-DFCP1 WT were treated with 2.5 µg/ml Puromycin for 2 h, fixed, stained with anti-p62 and anti-LC3B antibodies, and analyzed by super-resolution microscopy. Arrowheads denote omegasome rings with a lumen. Scale bar: 0.5 µm. (b) Cells treated as in (a) were stained with anti-p62 and anti-ubiquitin K63 antibodies and analyzed by super-resolution microscopy. Scale bar: 0.5 µm. (c) Parental U2OS, DFCP1 KO and KO rescue lines stably expressing mNG-DFCP1 WT, K193A or T189A were treated with 2.5 µg/ml Puromycin for 2 h, fixed, stained with anti-p62 antibody, and analyzed by confocal microscopy. Quantifications in Fig.4c. Scale bar: 10 µm. (d) Example of a mitophagy event. U2OS cells stably expressing mNG-DFCP1 WT and SNAP-mito were treated for 18 h with 0.5 mM DFP and imaged live. Note the lumen of the omegasome ring, through which a part of the mitochondria is threaded. Scale bar: 1 µm. Representative movie for > 10 events. (e) Knockdown validation for Fig.4e with Western Blot. Quantification of DFCP1- and ATG13 bands using Actin as a loading control. Values were normalized to control siRNA. Bars: ±SEM, 3 experiments. Statistics: One sample t-test, comparing against a theoretical mean of 100. (f) RPE-1 cells stably expressing mitochondrial mKeima probe and siRNA resistant mNG-DFCP1-WT, K193A, or T189A were depleted of endogenous DFCP1 by siRNA transfection for 2 d in complete medium (CM) before treatment with 50 nM Torin1 to induce mitophagy. Cells were harvested and analyzed by Flow Cytometry to determine the ratio of lysosome-localized mito-mKeima to free, cytoplasmic mito-mKeima. Values were normalized to the parental control. Bars: ± SEM, 3 experiments. Statistics: One-sample t-test, comparing against a theoretical mean of 100. (g) Knockdown validation for (f) with Western Blot. (h) U2OS parental cells, DFCP1-KO and KO rescue lines stably expressing mNG-DFCP1 WT, K193A or T189A were grown in complete medium, fixed, stained with Hoechst33342, and analyzed by confocal microscopy. Arrows point to micronuclei. Quantifications in Fig.4h. Scale bar: 20 µm. Same images as in Extended Data Fig.6b, which show p62 accumulation upon DFCP1 KO. Quantification of MN in Fig.4h. (i) Parental U2OS cells were depleted of endogenous DFCP1 by siRNA transfection for 2 d in complete medium, fixed, stained with Hoechst33342, and analyzed by confocal microscopy. Arrows point to micronuclei. Same images as in Extended Data Fig.6a, which show p62 accumulation upon DFCP1 depletion. (j) Quantification of number of micronuclei per cell from (i). Error bars denote mean ± SEM, 4 experiments. Each plotted point represents the mean value of one experiment. One-way ANOVA, Dunnett’s post hoc test: ** p < 0.01; **** p < 0.0001; ns: not significant. Cells analyzed per condition: NT (non-transfected): 575, siCtr: 507, si#1: 464, si#2: 504, si#3: 507. Re-analysis of the images from dataset Extended Data Fig.6a for scoring of micronuclei. (k) Quantification of number of micronuclei in parental A431 and DFCP1 KO cells. Cells have been grown in complete medium, fixed and stained with DAPI, and analysed by widefield microscopy. Two-sided Fisher’s exact test p < 0.0001. Data pooled from n=2 experiments. Cells analysed in total: WT=1230, KO=1148.

**Extended Data Fig.8:**
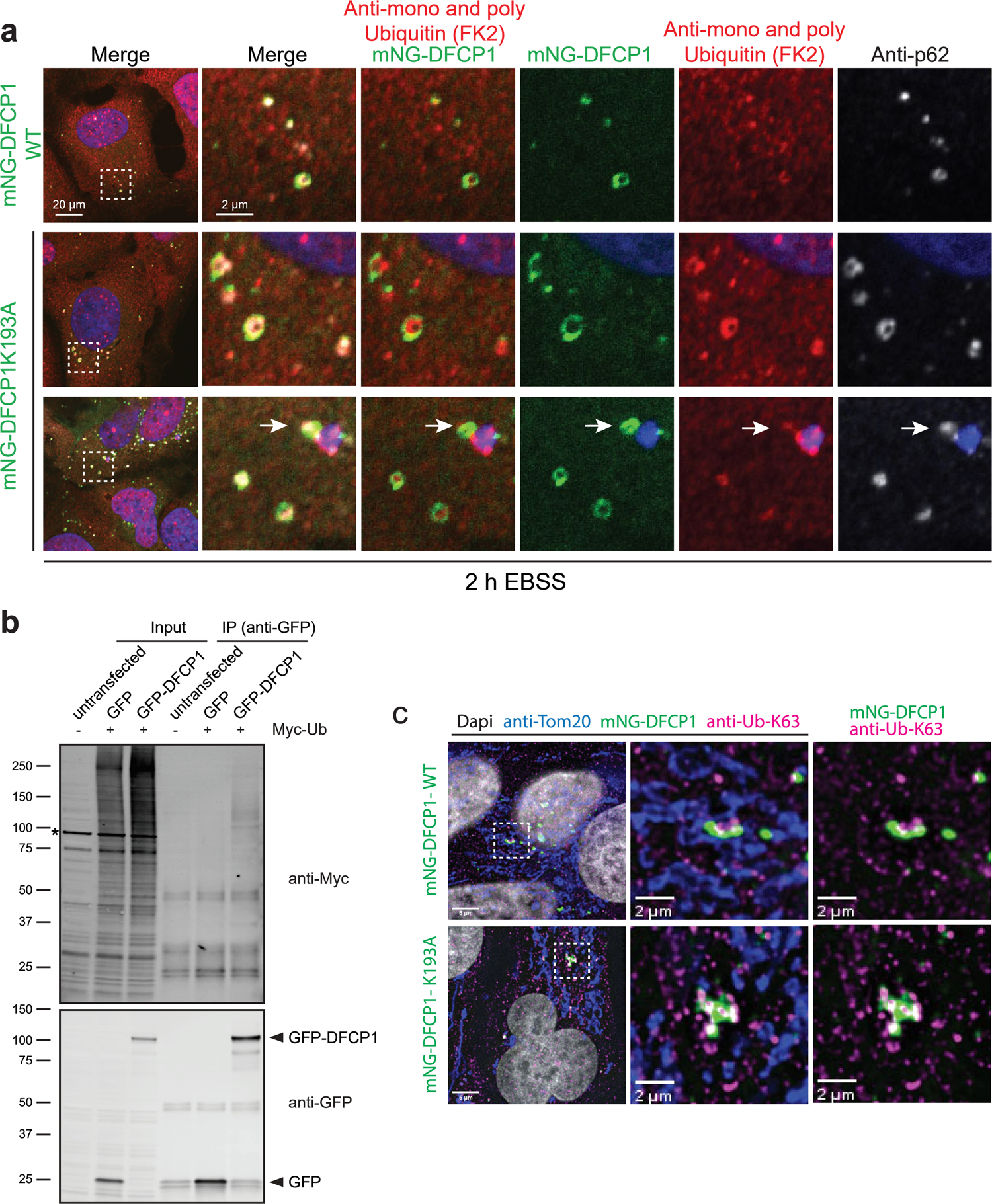
DFCP1 interacts with ubiquitin. (a) U2OS DFCP1 KO lines stably expressing DFCP1 WT or K193A were depleted of endogenous DFCP1 by siRNA, treated with EBSS for 2 h, fixed and stained with anti-mNG, anti-Ubiquitin and anti-p62 antibodies and analyzed by confocal microscopy. Arrows denote an omegasome localizing to a micronucleus. Scale bar 20 µm, insets 2 µm. (b) Co-IP of transfected HeLa cells. Western blot is representative for 3 independent experiments. Asterisk (*) indicates an unspecific band. (c) U2OS DFCP1 knockdown/rescue lines stably expressing mNG-DFCP1 WT or K193A were treated with DFP 0.5 mM for 18h to induce mitophagy, fixed, stained with anti-Tom20, anti-mNG and anti-Ub-K63 antibodies, and analyzed by widefield microscopy. Scale bar 5 µm, insets 1 µm.

**Extended Data Fig.9:**
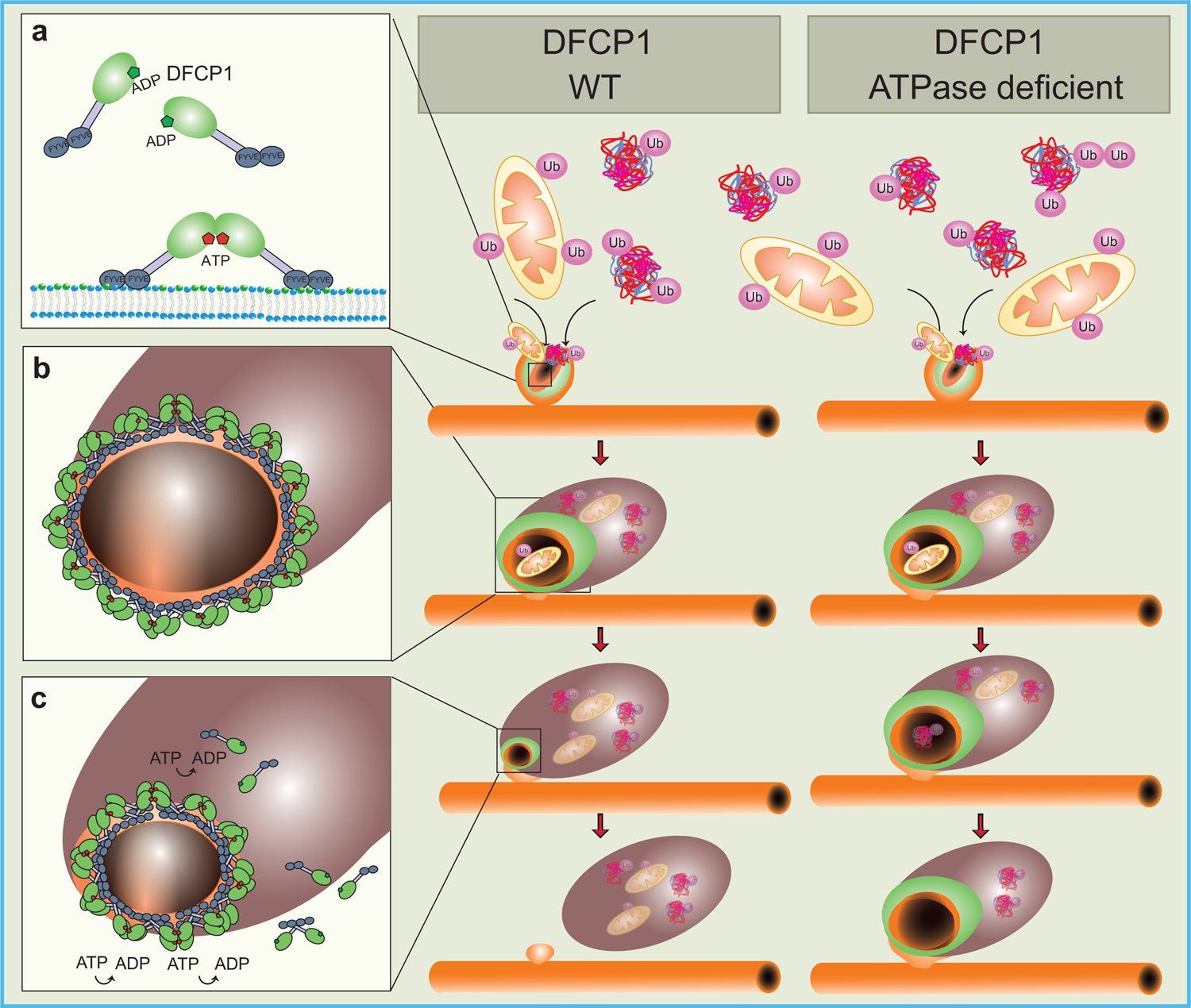
Model of how DFCP1 acts at the omegasome during selective autophagy. (a) Omegasome initiation: Upon induction of selective autophagy, PtdIns3P is generated at specialized subregions of the ER, at omegasomes. The FYVE domains of DFCP1 bind to these PtdIns3P pools and DFCP1 is assembled at the omegasome. ATP-binding of DFCP1 leads to its multimerization and could potentially generate a platform to bind ubiquitinated cargo. The phagophore is assembled within the omegasome cradle. (b) Omegasome maturation: the omegasome ring expands, accumulating DFCP1 dimers which form larger assemblies. This accumulation is likely driven by PtdIns3P at the omegasome. Nucleotide binding and multimerization stabilize DFCP1 at the omegasome. Omegasome constriction: Repeated cycles of ATP loading and hydrolysis drive the constriction of the omegasome ring. Constriction can either be achieved by transient disassembly of DFCP1 and the loss of some subunits, or alternatively, DFCP1 molecules can repeatedly engage neighboring molecules and constrict the membrane by a ratchet-like mechanism. DFCP1 mutants – either nucleotide binding defective of hydrolysis defective – cannot perform this repeated cycling. This leads to an inefficient constriction of the omegasome ring, the accumulation of omegasomes, and an inefficient degradation of selective cargo.

## Notes

### Competing Interest Statement

The authors have declared no competing interest.

